# Social Processing in the Amygdala: Single-Nucleus RNA-Sequencing Reveals Distinct Neuronal Responses to Dominant and Subordinate Cues

**DOI:** 10.1101/2025.11.24.690183

**Authors:** Madeleine F. Dwortz, Hans A. Hofmann, James P. Curley

## Abstract

The amygdala serves as a critical neural hub for interpreting social cues, with its distinct subregions and diverse neuronal populations playing specialized roles in processing these signals. This study employs single-nucleus RNA sequencing (snRNA-seq) to characterize the amygdala’s neuronal responses to olfactory cues associated with social dominance, uncovering distinct activation patterns within glutamatergic and GABAergic populations. We find that a glutamatergic cluster, characterized by expression patterns closely aligned with glutamatergic *Slc17a7* (VGLUT1) medial amygdala (MeA) neurons, preferentially responds to dominant cues. In contrast, a larger glutamatergic *Slc17a6* (VGLUT2) cluster associated with neurons of the MeA, as well as cortical and basomedial amygdala, exhibits a heightened response to subordinate cues, underscoring the MeA region’s role in processing social olfactory information. Additionally, a glutamatergic cluster resembling dorsal endopririform (EPd) neurons responded more to dominant stimuli, supporting the EPd’s role in olfactory perception. We also identified a GABAergic cluster with elevated dopamine receptor 2 (*Drd2*) expression that predominantly responds to dominant cues, consistent with this receptor’s known role in mediating threat responses. Through gene co-expression network analysis, we linked gene expression within neuronal clusters to specific biological processes. These findings reveal distinct neuronal and molecular mechanisms underlying social processing, particularly in response to dominant and subordinate olfactory signals, thus enhancing our understanding of the neural substrates of social behavior.

## Introduction

Social perception is a critical neural function for animals living in social groups. The brain’s ability to perceive social context crucially guides appropriate social behaviour, enabling animals to form and maintain strong social relationships (Adolphs 2001, O’Connell and Hofmann 2011, Dwortz et al. 2022). This is especially important in a dominance hierarchy, a common form of social organization, where the contextually appropriate expression of dominant or subordinate behaviour determines an individual’s access to resources and chances of survival. Decades of research show that the amygdala is crucial for proper social functioning in dominance hierarchies and that it may be especially important for the perception of dominance cues (Rosvold, Mirsky, and Pribram 1954, Adolphs 2003). For instance, activity in the primate basolateral amygdala (BLA) is positively associated with the dominance status of perceived individuals’ faces (Zink et al. 2008; Munuera, Rigotti, and Salzman 2018). In mice, amygdala subregions innervated by olfactory neurons, such as the posteroventral medial amygdala (MeApv), have been shown to increase their activity in response to dominant olfactory cues (W. Lee et al. 2021; Veyrac et al. 2011). Our recent work in mice suggests that similar response patterns extend to dorsal and anteroventral MeA subregions (MeAd, MeAav), BLA, and dorsal endopiriform nucleus (EPd) (Dwortz and Curley 2025). While many specific subregions within the amygdala contribute to processing social cues, the extensive heterogeneity of this brain region suggests that interconnected nuclei and diverse neural processes underpin social perception.

The amygdala is a highly heterogeneous structure, and recent advancements in single-cell profiling technologies have greatly enhanced our understanding and appreciation of its cellular diversity (Swanson and Petrovich 1998; Yu et al. 2023; Hochgerner et al. 2023). The amygdala contains a diverse array of neuronal cell types, including excitatory principal neurons and inhibitory interneurons. These cell types are intricately modulated by a complex network of neuromodulators, including dopamine (DA) and neuropeptides such as oxytocin (Oxt) and vasopressin (Avp), which play crucial roles in regulating social behaviours and emotional responses (de la Mora et al. 2010; Yao et al. 2017; Arakawa, Arakawa, and Deak 2010; Fadok, Dickerson, and Palmiter 2009; Lee, Lee, and Kim 2017). The expression of Oxt and Avp receptors is particularly prominent in the medial and central nuclei of the amygdala, areas that are key to processing social stimuli (Dębiec 2005; Ferguson et al. 2001; Ferretti et al. 2019). Moreover, the amygdala hosts a variety of projection neurons that connect different nuclei within the amygdala to other brain regions (Janak and Tye 2015; McGarry and Carter 2017; Kita and Kitai 1990; Dong, Petrovich, and Swanson 2001). Accounting for this cellular diversity is critical for elucidating how the amygdala processes social information.

In the present study, we captured the amygdala’s response to social dominance cues using single-nucleus RNA-sequencing (snRNA-seq). This method allowed us to simultaneously analyze responses of multiple amygdala subregions and characterize activated neurons at a granular, cell-type-specific level. Specifically, we exposed mid-ranking mice living in dominance hierarchies to urinary cues from familiar dominant and subordinate individuals and then used single-nucleus RNA-sequencing (snRNA-seq) to measure transcriptional responses of multiple amygdala subregions. To characterize activated neurons at a granular, cell-type-specific level we examined the expression of *Homer1*, an immediate early gene (IEG). IEGs are a class of genes that are rapidly and transiently transcribed upon cell membrane depolarization and, therefore, facilitate the identification of activated neurons. We further analyzed neurotransmitter gene expression and mapped neurons to specific amygdala sub-regions with the Allen Brain Cell database to aid in interpreting the functional implications of our findings. We hypothesized that exposure to dominant and subordinate social cues would elicit transcriptionally distinct patterns of neuronal activation across amygdala subregions. Specifically, we expected that dominant social cues would preferentially recruit excitatory neurons within the medial amygdala, reflecting specialized encoding of socially salient olfactory information. Our results confirm the heterogeneity of the amygdala, revealing a significant presence of both glutamatergic and GABAergic neurons, each distinguished by distinct subtypes. Specifically, we observed that glutamatergic VGLUT1 neurons whose transcriptional signatures were most closely aligned with MeA reference profiles exhibited heightened responses to the dominant olfactory cue. We also found that a cluster of glutamatergic VGLUT2 neurons with profiles matching those in the medial-cortical and basomedial amygdala regions showed a stronger response to the subordinate olfactory cue. This cluster notably co-expressed both *Oxt* and *Avp* preprohormones at relatively higher levels compared to other clusters. Additionally, we found that *Drd2*-expressing striatal-like GABAergic neurons predominantly responded to dominant olfactory cues. These results demonstrate that multiple mechanisms within the amygdala work together to process social cues.

## Methods

### Subjects and housing

We used 18 male CD-1 mice (*Mus musculus domesticus*) aged 7 to 8 weeks from Charles River Laboratory (Houston, TX, USA). Upon arrival, mice were marked for identification using blue non-toxic markers (Stoelting Co.) and housed in groups of three in standard cages with pine shaving bedding. For the duration of the experiment (25-26 days), subjects were housed in standard cages with pine shaving bedding under a reverse dark–light cycle and provided standard chow and water *ad libitum*. All procedures were conducted with approval from the University of Texas at Austin Institutional Animal Care and Use Committee (IACUC – Protocol No: AUP-2019-00338).

### Behavioural observations and dominance analysis

Groups were observed for one hour each day for the first five days of group housing and then no less than one hour every other day for the remainder of the study, between 11:00 and 17:00 during the dark phase of the reverse light cycle when mice were most active. During observation periods, observers used all-occurrence sampling to document fighting, chasing, mounting, submissive postures, and fleeing behaviours, noting winners and losers (see **Table S1** for ethogram). Behavioural data analysis was then conducted in R Studio using the Compete package (Curley 2016). The cumulative wins and losses for each individual over the housing period were compiled into frequency win/loss sociomatrices for each cohort (**Fig S1**). From these sociomatrices, we calculated the Directional Consistency (DC) of dominance interactions, and David’s Scores (DS) (de Vries 1995). DC assesses the degree to which all agonistic interactions in a group occur in the direction of the more dominant individual to the more subordinate individual within each relationship. It is equal to (H-L)/(H+L) where H is the frequency of behaviours occurring in the most frequent direction and L is the frequency of behaviours occurring in the least frequent direction within each relationship. The significance of DC values was evaluated using a randomization test (Leiva, Solanas, and Salafranca 2008). DS provides an individual dominance rating and ranking for each individual in a group, determining the overall success of each individual at winning contests relative to the success of all other individuals. Briefly, it is derived from the proportion of wins and losses of each individual and is corrected for the frequency of agonistic interactions (de Vries 1995). DS above 0 typically reflects that an animal is socially dominant whereas a DS below 0 typically indicates that an animal is socially subordinate. The most dominant and subordinate individuals in each hierarchy had the highest and lowest DS, respectively.

### Urine collection and odor exposures

Urine collection began one week prior to the odor presentations, starting on days 17 and 18 of group housing, and concluded one day before the odor presentations on days 24 and 25. Urine was collected from the most dominant and subordinate group members between 7 and 10 am, during the light cycle. Urine was collected by gently scruffing the mouse, applying a small amount of pressure on the bladder and collecting deposited urine directly into an Eppendorf tube. Samples were then immediately stored at –80° C.

On the morning of odor presentations, urine collected over the previous week was thawed on ice and then pooled into a single sample that remained at room temperature for approximately one hour. Odor presentations occurred between the hours of 11 am and 1 pm, during the dark phase of the light cycle. Mid-ranked subjects were first transferred to a separate behavioural testing room and placed in a testing chamber with a thin layer of fresh pine shaving bedding. They habituated to the chamber for 30 minutes and were then gently scruffed and the experimenter pipetted 15 µl of urine onto the nasal groove. Subjects were presented with urine samples from either the most dominant or subordinate member of their home-cage (n=). They were then placed back into the chamber, and the experimenter pipetted 1000 µl of urine onto the bedding. Subjects remained in the chamber for an additional 30 minutes and were sacrificed. Immediately upon sacrifice, brain tissue was flash-frozen in dry ice-cooled hexanes and stored at –80°C until further processing.

### Amygdala microdissection

Brain tissue was transferred to a −20° C cryostat chamber (Leica Microsystems), where it was embedded in OCT and incubated for approximately 30 minutes before amygdala dissection. We used the publicly available Allen Mouse Brain Atlas (Allen Reference Atlas – Mouse Brain, atlas.brain-map.org) to identify and section regions of interest. Four 300 µm slices were made onto glass slides, and the amygdala was dissected bilaterally using sterile surgical blades (approximately slides 65 to 77 of the Allen Mouse Brain Atlas). The amygdalae from each of the three mid-rank subjects per stimulus group (dominant and subordinate stimuli) were pooled into a single 0.2 ml RNAse-free tube and stored at –80°F until further processing.

### Nuclei dissociation, library preparation and sequencing

Samples were removed from the –80°C freezer and briefly brought to room temperature. We then isolated nuclei using a procedure adapted from Salem et al. 2024 (Salem et al. 2024). Samples were homogenized in 1.5 ml Nuclei EZ Lysis Buffer (Sigma # NUC101) with 0.2 U/ul RNAse inhibitor (NEB # ML314L) in pre-chilled 2mls KIMBLE Dounce tissue grinders (Sigma Aldrich D8938). Three up-and-down motions of the dounce sufficiently homogenized the samples. Lysate was then filtered through a 35-μm cell strainer and centrifuged at 900 rcf for 5 minutes at 4 °C. The pellet was resuspended in 700 µl of a 25% iodixanol mixture (2% BSA in 1X PBS supplemented with 0.2 U/ul RNAse inhibitor mixed with 60% iodixanol Optiprep medium, Sigma-Aldrich # D1556). This suspension was then layered on top of 700 µl of a 29% iodixanol cushion in a new 1.5 ml tube and centrifuged at 8,000 rcf for 30 minutes at 4°C. Supernatant was discarded and the pellet was resuspended in 150 μl buffer (2% BSA in 1X PBS supplemented with 0.2 U/ul RNAse inhibitor). This suspension was again filtered through a 35-μm cell strainer twice. The final eluent with nuclei was stained with DAPI and the concentration of nuclei was approximated using a Countess Automated Cell Counter (Thermo Fisher Scientific Single nuclei libraries were prepared by the Genomic Sequencing and Analysis Facility at UT Austin using the Chromium Next GEM Single Cell 3’ Kit v3.1 (10× Genomics, PN-1000128) with a target cell count of 10,000 cells. Libraries then were sequenced on a NovaSeq 6000 using an S4 flow cell.

### Initial snRNA-seq data processing

We used the Cell Ranger pipeline (10x Genomics) to align raw sequencing data to the mouse genome (GRCm39, Ensembl release 2024-A), perform cell calling, and generate gene-by-cell UMI count matrices. These matrices were then imported into R (RStudio, v4.3.1) and processed using Seurat (v4.3.0.1) (Hao et al. 2024; 2021; Stuart et al. 2019; Butler et al. 2018). We used the Souporcell pipeline (Heaton et al. 2020) to demultiplex each sample pool and identify three putative individuals (**Fig S2A,B**). We then filtered each dataset to ensure comparable cell count and quality between samples. Souporcell also aided in the detection and removal of doublets. (**Fig S2C**). Poor-quality nuclei were identified based on mitochondrial DNA content and MALAT1 expression. Using MALAT1, a nuclear-retained non-coding RNA universally expressed across cell types, as a quality control metric is a recommended approach in processing single-cell and single-nucleus data, as low MALAT1 expression is indicative of droplets lacking nuclei (Clarke and Bader 2024). We retained cells with 500–7,000 expressed genes, less than 1% mitochondrial DNA, and log normalized MALAT1 expression greater than 5 (**Fig S3**). Gene counts were normalized with Seurat’s NormalizeData function, which divides by total expression, scales to a common factor (10,000), and applies log transformation (Stuart et al. 2019).

### Data integration, clustering and annotation

We integrated the datasets derived from dominant– and subordinate-urine exposed subjects to identify common cell-type clusters using Seurat (**Fig S5**). This integration reduced batch effects and enabled direct comparison of shared cell-type clusters across conditions, facilitating the identification of cellular heterogeneity and accurate cell-type annotation. To ensure that clustering was based on cell-type similarity rather than activation states, we first removed IEGs. Based on existing literature, the following IEGs were excluded: *Btg2, Jun, Egr4, Fosb, Junb, Gadd45g, Fos, Arc, Nr4a1, Npas4, Coq10b, Tns1, Per2, Ptgs2, Rnd3, Tnfaip6, Srxn1, Tiparp, Ccnl1, Mcl1, Dnajb5, Nr4a3, Fosl2, Nptx2, Rasl11a, Mest, Sertad1, Egr2, Midn, Gadd45b, Dusp6, Irs2, Plat, Ier2, Rrad, Tpbg, Csrnp1, Peli1, Per1, Kdm6b, Inhba, Plk2, Ifrd1, Baz1a, Trib1, Pim3, Lrrk2, Dusp1, Cdkn1a, Pim1, Sik1, Frat2, Dusp5, Egr1, Homer1, Bdnf, Nr4a2* (Hochgerner et al. 2023). Next, a set of 2,000 highly variable features was selected to identify consistent patterns of variation across datasets using Seurat’s FindVariableFeatures function. We then computed PCA projections, determining the number of principal components based on the point where the explained variance reached an optimal balance. Seurat’s FindNeighbors function was then used to calculate the nearest neighbors for each cell, followed by the FindClusters function to apply the Louvain algorithm and identify clusters of transcriptionally similar cells (**Fig 1A**).

**Fig 1.**
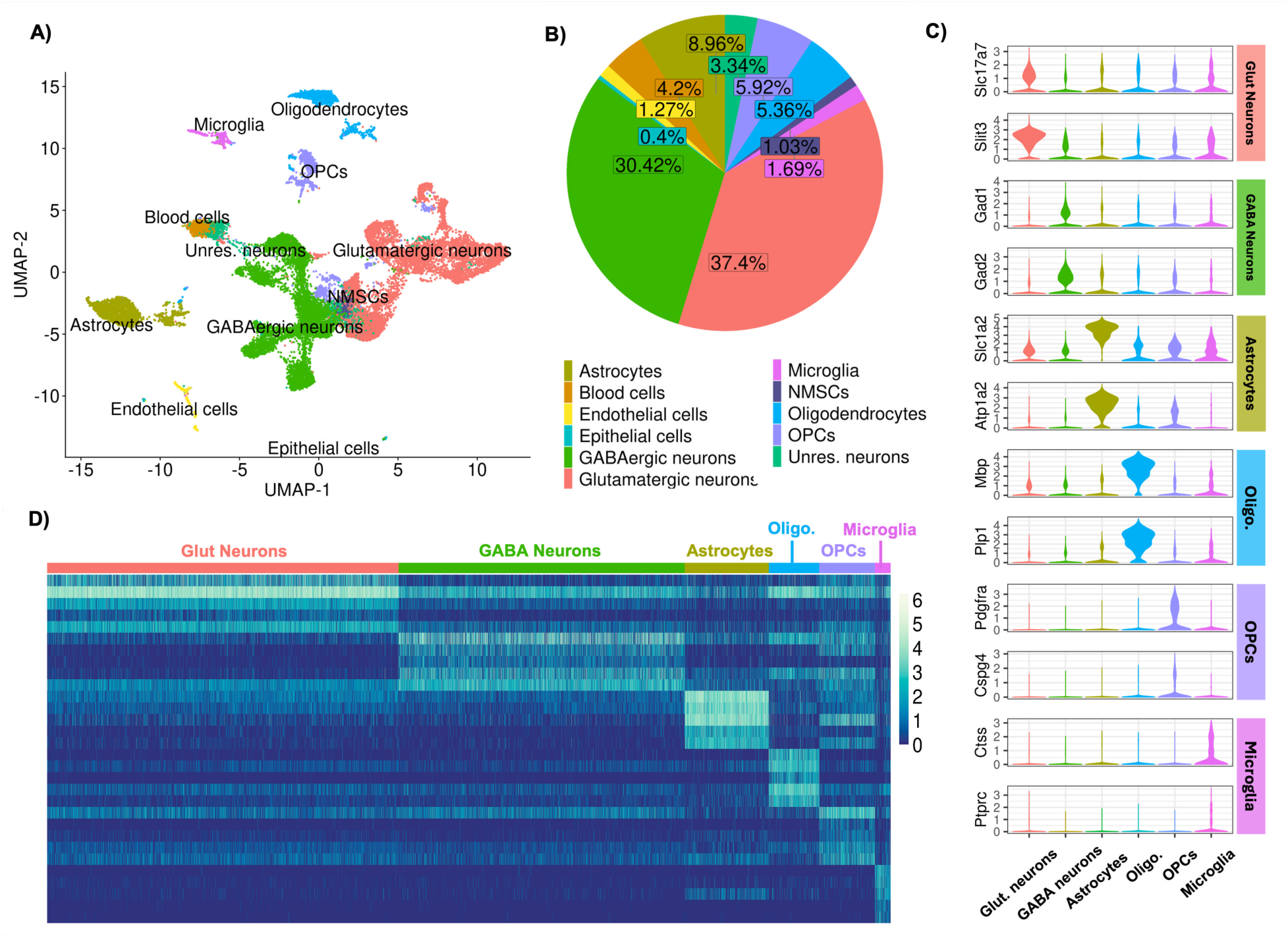
Cell Type Diversity in the Amygdala. A) Uniform Manifold Approximation and Projection (UMAP) plot showing the clustering of 22,695 nuclei in the integrated dataset. Nuclei are color-coded by identified cell types, including astrocytes, blood nuclei, endothelial nuclei, epithelial nuclei, GABAergic neurons, glutamatergic neurons, microglia, oligodendrocytes, oligodendrocyte progenitor nuclei (OPCs), and uncharacterized neurons. Labels reflecting the predominant cell type within each group. This visualization highlights the clustering of transcriptionally similar nuclei by their assigned cell type. B) Pie chart showing the relative abundance of each cell type. Glutamatergic neurons (37.4%) and GABAergic neurons (30.42%) represent the most prevalent types. C) Violin plots display the distribution of expression levels for canonical markers specific to each cell type, supporting nuclei classifications. D) Heatmap showing the expression levels of top marker genes for various cell types, selected based on an average Log2(Fold Change) > 1 and a Bonferonni-adjusted P value < 0.05, across individual nuclei. Normalized Log expression levels are scaled from low (blue) to high (light yellow).

We assigned cell types to the identified clusters using multiple annotation approaches. First, Seurat’s FindAllMarkers function was used to identify cluster-specific marker genes. This function uses a Wilcoxon Rank Sum test to compare gene expression levels between each cluster and all other cells and provides Bonferroni-corrected p-values. In particular, this analysis revealed a distinct cluster of blood cells characterized by high expression of *Ttr*, *Hbb-bs*, and *Hba-a1* (Hochgerner et al. 2023). To perform reference-based classification, we applied SingleR with reference datasets from Celldex (Aran et al. 2019) to assign broad cell classes including neurons, astrocytes, oligodendrocytes, endothelial cells, and epithelial cells. We next used ScType (Ianevski et al. 2022) to refine annotations of specific subtypes, such as oligodendrocyte precursor cells (OPCs), based on curated marker databases. Finally, we employed the Allen Brain Cell “Map My Cells” (MMC) tool to achieve finer anatomical resolution and facilitate comparability with future datasets. MMC uses a hierarchical reference-mapping algorithm based on the Allen Brain Atlas to assign nuclei to specific cell types and returns a bootstrapped probability score reflecting assignment confidence (Z. Yao et al. 2023).

We examined the results of these various annotation methods in the 10x Loupe Browser (10x Genomics) and assigned cells based on agreement across methods. Cells with conflicting annotations were re-clustered using the full integration and clustering pipeline, enabling the selection of more biologically relevant genes for feature selection. We next analyzed glutamatergic and GABAergic neurons separately, re-running the clustering workflow within each group to resolve finer neuronal subtypes. To identify overexpressed genes associated with each cluster, we again employed Seurat’s FindAllMarkers function.

### Identifying differentially activated neuronal populations

We aimed to identify differences in neuronal activity between dominant and subordinate stimuli by comparing the proportions of *Homer1-*expressing (*Homer1⁺*) neurons within each glutamatergic and GABAergic cluster for each individual mouse genotype from the sample pools. We used binomial generalized linear mixed-effect models (GLMMs), incorporating the total number of cells per cluster per individual as weights to account for varying cell counts. To explore the functional characteristics of differentially ‘activated’ cell populations, we analyzed the MMC subclass annotations assigned to each neuronal cluster. We also used a candidate gene approach and examined neurotransmitter gene expression across subclusters. Additionally, we performed differential gene expression analysis between *Homer1⁺*and *Homer1⁻*neurons using Seurat’s FindAllMarkers function and Gene Ontology (GO) Enrichment analysis to confirm that processes related to cell activation were activated in these nuclei.

### Weighted Gene Co-expression Network Analysis

We also conducted a weighted gene co-expression network analysis for high dimensional data (hdWGCNA) using the hdWGCNA R package to identify modules of co-expressed genes across the integrated datasets for glutamatergic and GABAergic nuclei (Morabito et al. 2023). We selected soft power thresholds of 12 and 10 for glutamatergic and GABAergic nuclei, respectively, and only included genes expressed in at least 5% of nuclei in each cell type. We computed module eigengenes (MEs) using hdWGCNA’s ModuleEigengenes function. Within this function, we adjusted for potential sample quality and technical biases by regressing out the influence of total counts per cell, the percentage of mitochondrial genes expressed, and *Malat1* expression. GO enrichment analysis was then conducted using the Enrichr package to determine active biological pathways associated with each module. For each module, the corresponding gene list was queried against the GO:Biological Process, GO:Cellular Component and GO:Molecular Function databases. (Chen et al. 2013). We examined the expression of MEs and pathway activity across neuronal clusters and analyzed the overlap between cluster-specific gene markers and module-associated genes.

## Results

### Dominance

All six social groups of male mice established dominance hierarchies with a mean Directional Consistency (DC) of 0.752 ± 0.198 (**Fig S1**; **Table S2**). The stability and strength of each dominance relationship determined how subjects were assigned a dominant or subordinate stimulus. For instance, in cohort C, the mid-ranking mouse defeated the most dominant individual in nine interactions, resulting in a low DC score of 0.443. However, this mouse consistently defeated the most subordinate individual, indicating a stable dominance relationship with that opponent. Consequently, the mid-ranking mouse in cohort C received urine from the subordinate animal (see **Fig S1** for stimulus assignments and **Table S2** for full dominance results).

### Neuronal diversity within the amygdala

We identified 22,695 nuclei that passed the quality control filters (11,597 cells in the dominant-stimulus sample and 11,098 in the subordinate-stimulus sample) containing 125,267,275 total reads and 33,696 unique genes. These nuclei included neurons, astrocytes, oligodendrocyte precursor cells (OPCs), oligodendrocytes, blood cells, endothelial cells, non-myelinating Schwann cells (NMSCs), and epithelial cells (**Fig 1**; see **Table S3** for list of cell type gene markers). Neurons clustered robustly into glutamatergic (8,489 cells) and GABAergic (6,904 cells) clusters (**Fig 1A**). Ultimately, only a small fraction of unresolved neurons could not confidently be assigned to glutamatergic or GABAergic classes (**Fig 1B**). Glutamatergic and GABAergic populations were further clustered, and we identified 16 glutamatergic cell clusters (**Fig 2A-D**) and 11 GABAergic cell clusters (**Fig 2E-H**; see **Tables S4 and Table S5** for lists of neuronal cluster gene markers). Across neuronal clusters, there was some variability in the relative contributions of each genotype by random chance (**Fig S5C,F**).

**Fig 2.**
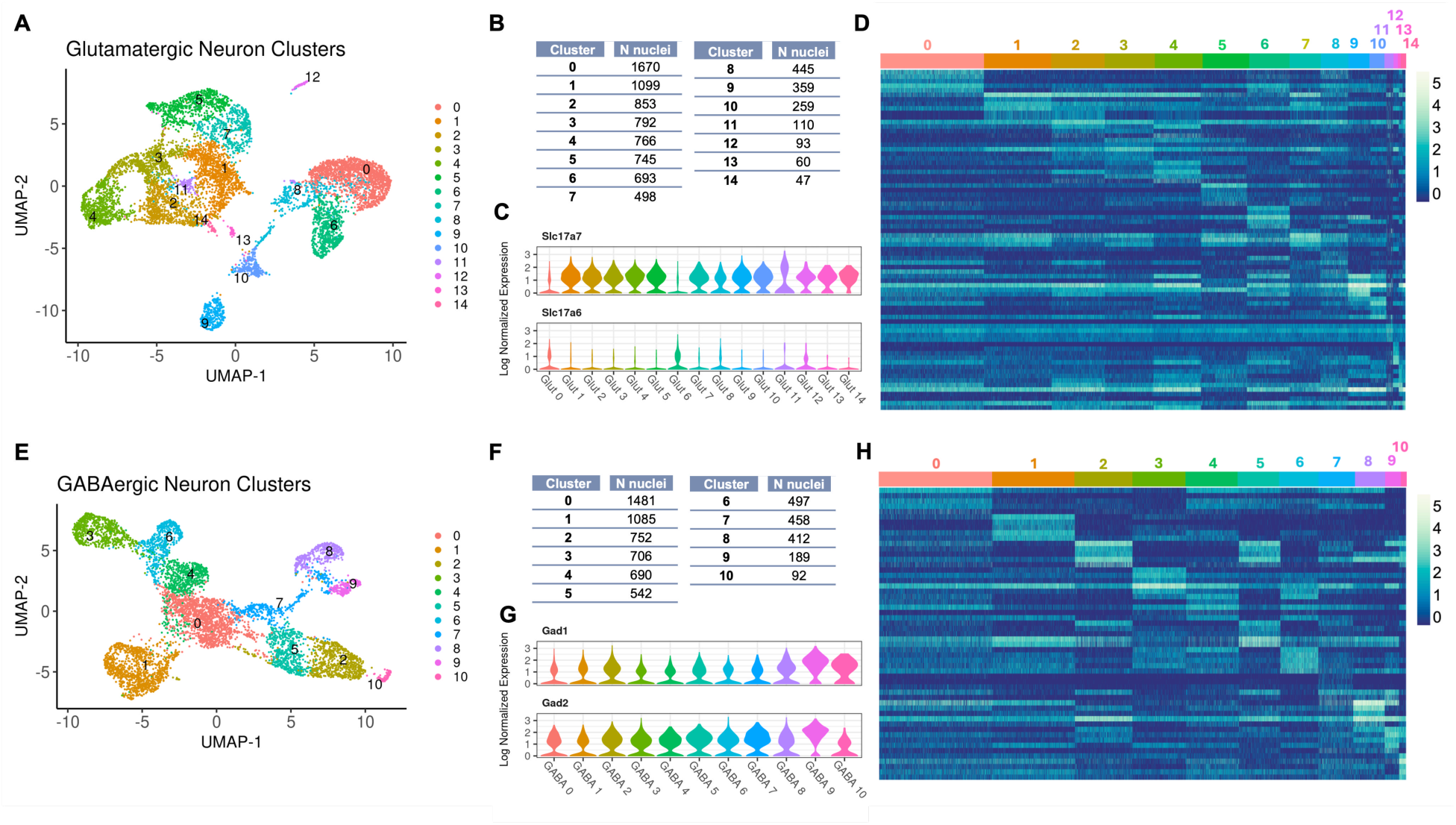
Clustering and Gene Expression Profiles of Glutamatergic and GABAergic Neurons. A, E) UMAP visualizations of glutamatergic (A) and GABAergic (E) neuron clusters. Nuclei are color-coded by cluster identity, with each color representing one of the 15 clusters for glutamatergic neurons and 10 clusters for GABAergic neurons. The clusters represent distinct subpopulations based on gene expression similarities. B, F) Tables showing the number of nuclei per cluster for glutamatergic (B) and GABAergic (F) neurons, respectively. Each cluster is assigned a unique identifier with the corresponding count of nuclei, providing a quantitative view of the distribution across clusters. C, G) Violin plots depicting the normalized expression levels of marker genes across clusters for glutamatergic (*Slc17a7* and *Slc17a6*) and GABAergic (*Gad1* and *Gad2*) neurons. D, H) Heatmaps showing the normalized expression of top marker genes for glutamatergic (D) and GABAergic (H) clusters. Rows represent genes selected based on an average Log2(Fold Change) > 1 and a Bonferonni-adjusted P value < 0.05. Columns are nuclei, grouped by cluster. Color intensity indicates the expression level, ranging from low (dark blue) to high (light yellow). These heatmaps provide a comprehensive overview of the gene expression landscape, highlighting genetic differences between the identified clusters.

We observed cluster-specific expression of *Slc17a7* (VGLUT1) and *Slc17a6* (VGLUT2) subtypes among glutamatergic clusters (**Fig 2C**). Specifically, VGLUT2 neurons were more prominent than VGLUT1 in glutamatergic clusters 0 and 6. In contrast, expression of GABAergic marker genes *Gad1* and *Gad2* were uniformly expressed across GABAergic clusters (GABAergic marker gene *Slc32a1* was very lowly expressed in nuclei) (**Fig 2G**).

We also examined the expression of neurotransmitter and receptor genes implicated in social processing across glutamatergic and GABAergic clusters (**Fig 3A,B; Table S6**). *Oxt* and *Avp* preprohormones were expressed across various subclusters (average log normalized expression in glutamatergic nuclei: *Oxt*: 0.074<u>0</u> ± 0.281, *Avp*:0.218 ± 0.467; GABAergic nuclei: *Oxt*: 0.0798 ± 0.305, *Avp*: 0.223 ± 0.484). In contrast, the expression of neuropeptide receptor genes was generally low across all nuclei, which was not unexpected, given that many neuropeptide GPCR mRNAs are localized to distal processes rather than the nucleus or soma, making them less readily detected in snRNA-seq (Cajigas et al. 2012) (average log normalized expression in glutamatergic nuclei: *Oxtr*: 0.0181 ± 0.130, *Avpr1a*: 0.000 ± 0.015, *Avpr1b*: 0.000415 ± 0.0181; GABAergic nuclei: *Oxtr*: 0.0205 ± 0.148, *Avpr1a*: 0.001 ± 0.035, *Avpr1b*: 0.000435 ± 0.0211). However, dopamine receptor genes were detectable (average log normalized expression in glutamatergic nuclei: *Drd1*: 0.040 ± 0.19, *Drd2*: 0.024<u>8</u> ± 0.166; GABAergic nuclei: *Drd1*: 0.055 ± 0.25, *Drd2*:0.0837 ± 0.329), GABAergic cluster 3 exhibited especially prominent *Drd2* expression compared to other clusters (average log normalized expression: 0.523 ± 0.720) (**Fig 3B**, **Table S5**).

**Fig 3.**
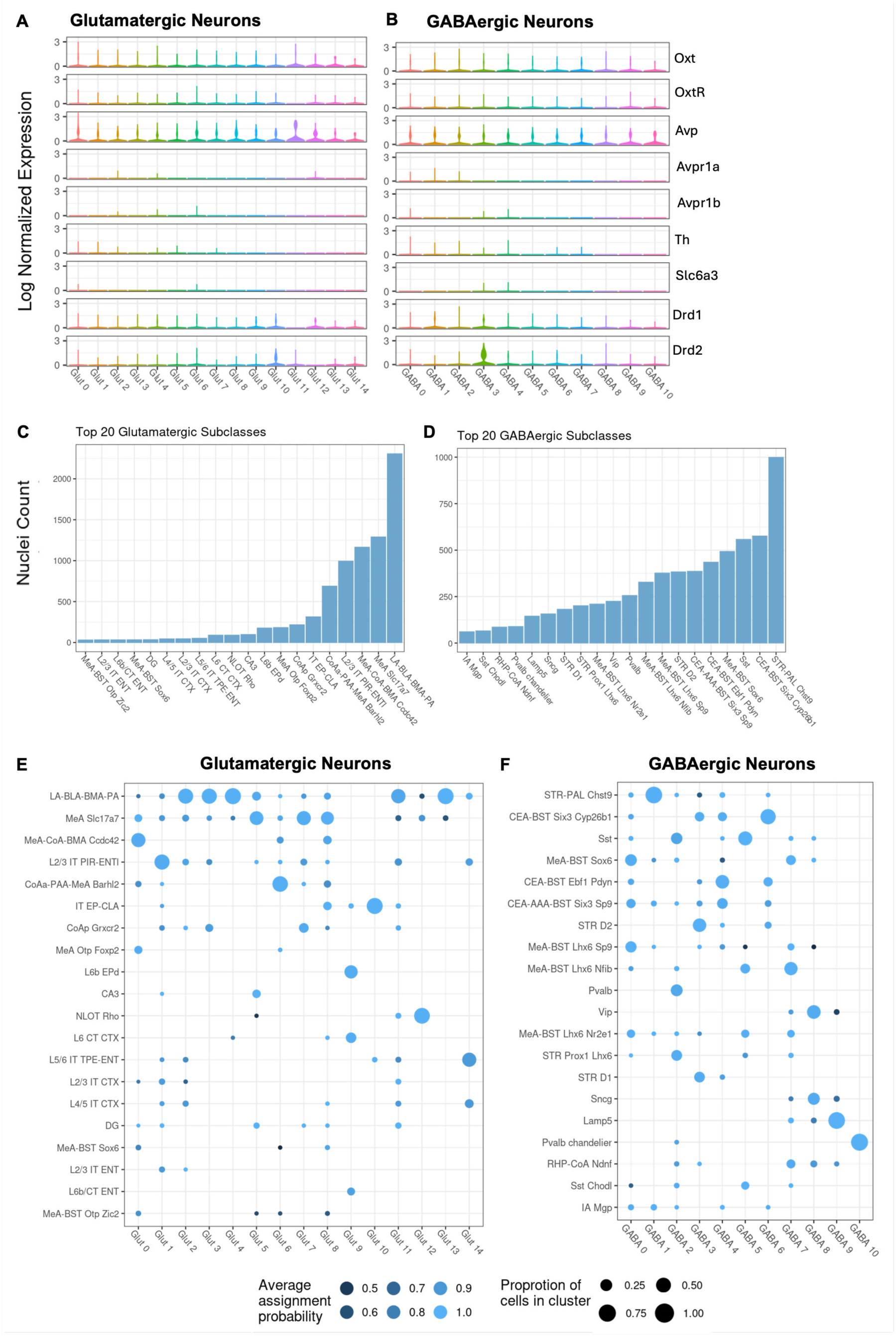
Characterizing Glutamatergic and GABAergic Clusters. A, B) Violin plots displaying the log-normalized expression of neurotransmitter genes that have been implicated in social processing across clusters within glutamatergic (A) and GABAergic (B) neurons. Each violin plot represents the distribution of expression levels for a specific gene, providing insights into the variation and expression levels that characterize each neuronal subtype (see Table S3.5 for a statistical summary of these expression levels). C, D) Bar graphs illustrating the counts of nuclei within the top 20 most populous subclasses as identified by the Map My Cells (MMC) method for glutamatergic (C) and GABAergic (D) neurons. These top 20 subclasses account for 95.69% and 91.03% of all glutamatergic and GABAergic neurons, respectively. E, F) Dot plots showing the distribution of neuronal subtypes across the glutamatergic (E) and GABAergic (F) neuron clusters. Dot size indicates the proportion of nuclei in each cluster assigned to the subtype and color intensity reflects the average assignment probability.

To assess the relative prominence of each subtype within clusters we further analyzed the distribution of neurons across distinct neuroanatomical subtypes using the MMC tool (**Fig 3C-F**). Clusters showed enrichment for specific MMC-derived assignments, which were annotated according to their reference brain region and characteristic gene markers. These assignments suggest that the transcriptional profiles of nuclei in our dataset closely resemble those cataloged in the Allen Brain Cell database. While definitive classification of these nuclei cannot be established, their expression patterns indicate strong similarity to known cell types. Most glutamatergic nuclei resembled cells originating from the centralized collection of amygdala nuclei consisting of the basolateral (BLA), basomedial (BMA), lateral (LA) and piriform (PA) amygdala: LA-BLA-BMA-PA. The next most prevalent sources of glutamatergic neurons were the medial (MeA), anterior cortical (CoAa), piriform (PAA) amygdala area, and piriform-entorhinal (PIR-ENT) neurons: MeA *Slc17a7*, MeA-CoA-BMA *Ccdc42*, L2/2 PIR-ENT, CoAa-PAA-MeA *Barhl2* (**Fig 3C**).

GABAergic neurons appeared to be comprised of two primary types: striatal or striatal-like neurons and local inhibitory neurons. The largest proportion was identified as striatal pallidal neurons (STR-PAL Chst9). Prominent sources of GABAergic nuclei also included the central (CEA), medial and anterior (AAA) amygdala and bed nuclei of the striatal terminalis (BST): (CEA-BST Six3 Cyp26b1, CEA-BST Ebf1 Pdyn, CEA-AAA-BST Six3 Sp9). Somatostatin-positive neurons (SST) were also prevalent (**Fig 3D**).

### Differential activation across neuronal clusters

As expected, *Homer1* was the most highly expressed IEG in glutamatergic and GABAergic neurons (Fig S6A,B) and it showed strong correlations with longer-timescale IEGs such as *Bdnf* (Tao et al., 1998) (**Fig S6C,D**). *Homer1* expression was generally higher among glutamatergic neurons, with the highest expression in glutamatergic cluster 2 (Glut 2) (mean = 1.11 ± 0.84) (**Fig 4A**). Among GABAergic neurons, cluster 3 (GABA 3) exhibited the highest expression (mean = 0.778 ± 0.747) (**Fig 4B**).

**Fig 4.**
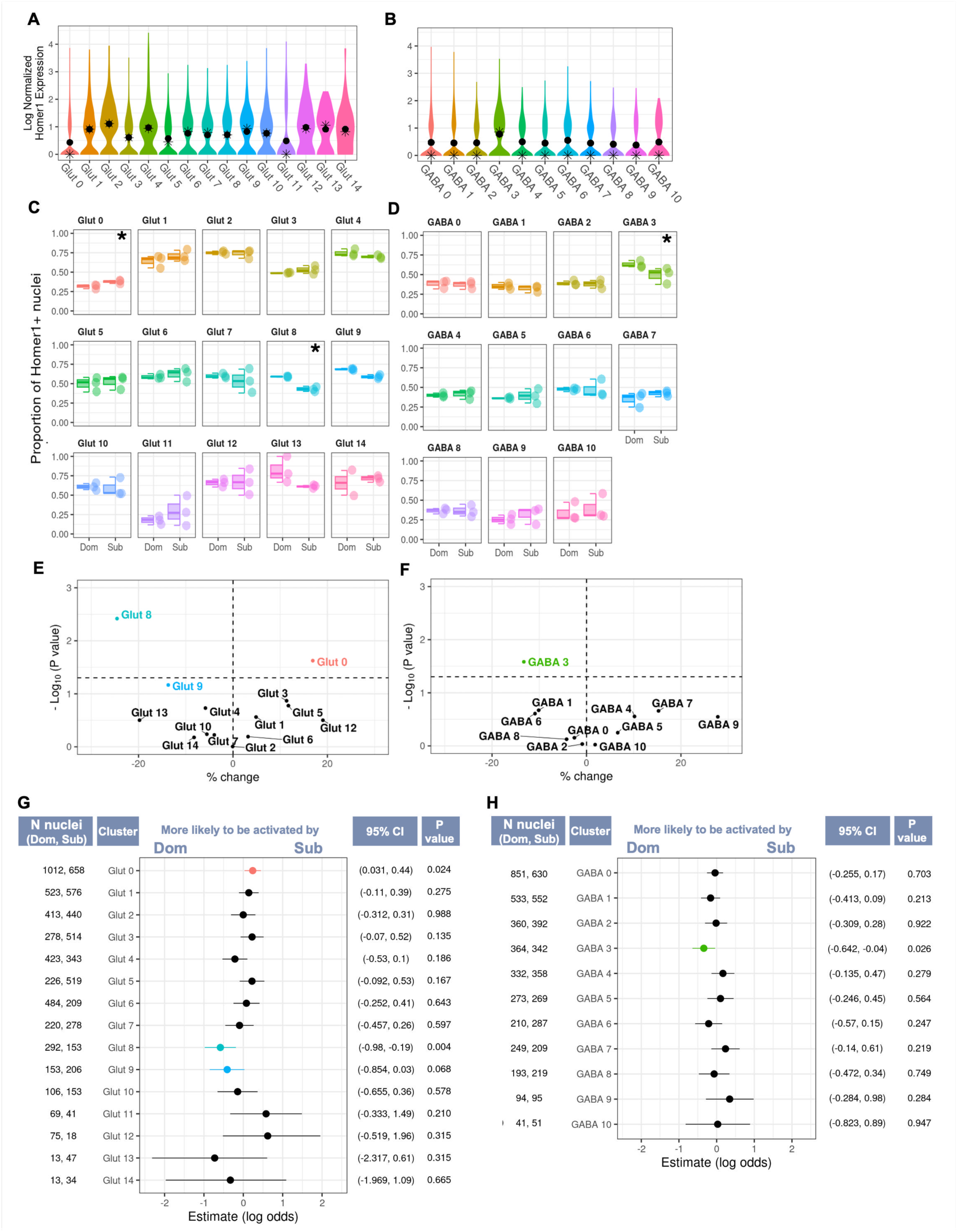
Analysis of Immediate Early Gene (IEG) Expression Across Glutamatergic and GABAergic Neuron Clusters. A, B) Violin plots show the distribution of *Homer1* expression across glutamatergic (A) and GABAergic (B) neuron clusters. The mean and medial expression level within each plot is marked by a dot and a star, respectively. C, D) Box plots show the proportion of *Homer1⁺* nuclei in response to dominant (Dom) and subordinate (Sub) stimuli across glutamatergic (C) and GABAergic (D) neuron clusters. The significance level of each comparison is indicated in the upper right corner (p < 0.05 (*), p < 0.01 (**), p < 0.001 (***), 0.05 < p < 0.07 (†). E, F) Volcano plots depict the percentage change in the number of *Homer1⁺* nuclei between Dom and Sub stimuli for glutamatergic (E) and GABAergic (F) neuron clusters. Data points represent individual clusters, with the x-axis showing the percentage change and the y-axis indicating the –log10(P value). Points to the right of the vertical dashed line signify significant increases, while those to the left signify decreases. G, H) Forest plots display the log odds of *Homer1* activation in response to Dom versus Sub stimuli for glutamatergic (G) and GABAergic (H) neuron clusters. Each point on the plot corresponds to the estimate for each cluster, with the horizontal lines representing 95% confidence intervals. Confidence intervals are specified to the right of the plot, followed by the P values. The number of nuclei in each cluster for Dom and Sub samples, the denominators used to calculate *Homer1⁺* proportions, are shown to the left of the plot.

To confirm that *Homer1* expression reliably marks neuronal activation and to identify additional genes associated with this active state, we compared transcriptional profiles of *Homer1*⁺ and *Homer1*⁻ nuclei (see **Table S9** for full DEG results). We found overexpression of phosphodiesterase (PDE) genes in glutamatergic and GABAergic *Homer1⁺* cells (**Fig S7A-D**). PDE’s control intracellular signaling by catalyzing second messenger cyclic nucleotides, which in turn control receptor-effector coupling, protein-kinase cascades, and transmembrane signal transduction. GO enrichment analysis of these overexpressed genes revealed terms related to “cyclic-nucleotide-mediated signaling” (see **Table S10** for GO analysis results). We specifically observed overexpression of *PDE10a* in both glutamatergic and GABAergic *Homer1⁺* neurons, along with Pde4b and Pde7b in glutamatergic and GABAergic *Homer1⁺* neurons, respectively. Additional GO enrichment terms associated with *Homer1⁺* neurons also underscored the processes related to post-synaptic activation, such as “regulation of postsynaptic membrane neurotransmitter receptor levels” and “protein localization to postsynaptic membrane”. This is consistent with the established role of *Homer1⁺*in interacting with excitatory metabotropic glutamate receptors at the postsynaptic density (Xiao et al. 1998; Vazdarjanova et al. 2002). In contrast, genes overexpressed in *Homer1⁻* nuclei were associated with GO terms related to cell homeostasis, such as “maintenance of location in cell” and “regulation of cellular component size”.

We applied general linear mixed-effect models (GLMMs) for binomially distributed data to proportions of *Homer1⁺* nuclei within each individual mouse genotype and neuronal cluster to determine if there was an association with the social status of the olfactory stimulus. We observed significant differences in the proportion of *Homer1⁺* nuclei in several glutamatergic and GABAergic clusters (**Fig 4C-H**; see **Table S8** for full GLMM results).

Glutamatergic cluster 8 (Glut 8) exhibited significantly greater proportions of *Homer1⁺*nuclei in response to dominant stimuli compared subordinate stimuli (Sub – Dom: β = –0.583 ± 0.202, P = 0.004, 95% CI: –0.980, –0.190). Notably, this cluster was enriched for excitatory MeA neurons. Within this cluster, 54.8% of nuclei were confidently identified as MeA *Slc17a7* neurons with a 99.0% average probability of assignment accuracy (avg. probability), while MeA-CA-BMA *Ccdc42* neurons constituted 14.3% (avg. probability = 95.6%) (**Fig 3E**; see **Table S7** for full MMC assignments). Additionally, proportions of *Homer1⁺* nuclei in glutamatergic cluster 9 (Glut 9) trended higher in response to dominant compared to subordinate stimuli (Sub – Dom: β =-0.410 ± 0.225, P = 0.068, 95% CI: –0.854, 0.028). Glut 9 was the only cluster enriched for dorsal endopiriform neurons (EPd) at 58.6% (avg. probability = 98.2%) (**Fig 3E**).

In contrast, glutamatergic cluster 0 (Glut 0) exhibited significantly greater proportions of *Homer1⁺* nuclei in response to subordinate stimuli compared to dominant stimuli (Sub – Dom: β = 0.238 ± 0.105, P = 0.024, 95% CI:0.031,0.444). Cluster Glut 0 was predominantly composed of excitatory MeA-CA-BMA *Ccdc42* neurons, accounting for 70.5% (avg. probability = 97.8%). (**Fig 3E, Table S7**). Notably, this cluster exhibited elevated expression of *Oxt* and *Avp* preprohormone transcripts compared to other neuronal clusters. Specifically, it showed the highest *Oxt* preprohormone expression and the second highest *Avp* preprohormone expression (average log normalized expression of *Oxt*: 0.110 ± 0.363; *Avp*: 0.316 ± 0.602; **Fig 3A, Table S6**).

We also observed stimulus-associated *Homer1⁺* expression among GABAergic neurons. GABA 3 exhibited significantly greater proportions of *Homer1⁺* nuclei in response to dominant compared to subordinate stimuli (Sub – Dom: β = –0.341 ± 0.153, P = 0.026, 95% CI: –0.642, – 0.041). This cluster also exhibited robustly higher *Drd2* expression compared to all other neuronal clusters (**Fig 3B**). In line with this observation, this cluster was enriched for striatal DRD2-expressing neurons (STR D2) based on the MMC analysis (**Fig 3F**).

To better understand and characterize the amygdala’s complex neuronal populations, we used hdWGCNA to identify and analyze modules of co-expressed genes distributed across neuronal clusters. Additionally, we performed Gene Ontology (GO) enrichment analysis on these modules, uncovering active biological processes in these clusters. We discovered five modules within glutamatergic neurons and seven within GABAergic neurons (**Fig S8, Fig S9**; see **Table S11** and **Table S12** for module gene assignments). We focused specifically on clusters exhibiting stimulus-associated *Homer1⁺* expression. Glut 0, which showed a higher proportion of *Homer1⁺*nuclei in response to subordinate stimuli, exhibited elevated expression of the brown module eigengene (ME) (**Fig S9A,B**). This was also supported by a significant overlap between its marker genes and those of the brown module (**Fig S9G**). GO terms associated with this module included terms included “chloride transport” and “regulation of postsynaptic membrane potential” (**Fig S9E**; see **Table S13** for GO analysis results). In response to dominant stimuli, Glut 9 exhibited a higher number of *Homer1⁺* nuclei and showed elevated expression of the green module, the smallest module composed of only 28 genes (**Fig S9B, Fig S9G**). GO terms associated with the green module were related to “cell-cell adhesion” and “cell junction assembly” (**Fig S9E**). Glut 8 did not exhibit elevated expression of any particular module, although it did share some marker genes with the brown and green modules (**Fig S9G**). GABA 3, which showed a higher proportion of *Homer1⁺* nuclei in response to dominant stimuli, exhibited a high degree of overlapping marker genes with the blue module and elevated expression of the blue ME (**Fig S9C,D,G**). The top GO terms for the blue module included terms related to cell activation, including “regulation of cation channel activity”, “chemical synaptic transmission” and “protein phosphorylation” (**Fig S9F**).

## Discussion

The amygdala serves as a central hub for processing social information and encompasses diverse subregions and neuronal types (Janak and Tye 2015, Li et al. 2017, Adolphs 2003, Hochgerner et al. 2023,). In the present study, we employed snRNA-seq to identify specific neuronal populations within the amygdala activated by social olfactory cues. Our analysis revealed distinct clusters of glutamatergic and GABAergic nuclei, each characterized by neurotransmitter systems and enrichment for specific neuroanatomical cell types. The heterogeneity found in our dataset largely corroborates existing amygdala taxonomy revealed by single-cell RNA-seq studies (Hochgerner et al. 2023). This neuronal diversity highlights the need for cell-type-specific analyses of social processing in the brain.

We identified several neuronal clusters that were differentially activated, as indicated by the expression of *Homer1*, by dominant and subordinate olfactory cues. Glut 8, enriched for MeA *Slc17a7* (VGLUT1) neurons, responded more to dominant stimuli. This finding aligns with our previous work showing that dominant urinary cues elevate c-fos immunoreactivity in the MeApv, a region with a high concentration of glutamatergic neurons (Keshavarzi et al. 2014; W. Lee et al. 2021). The MeA plays a critical role in processing social olfactory cues, acting as a pivotal relay station within the accessory or vomeronasal olfactory pathway, which primarily processes pheromonal cues (Raam and Hong 2021; Kevetter and Winans 1981; Dulac and Wagner 2006). Numerous studies have shown that activity within the MeA responds to socially-relevant olfactory cues (Li et al. 2017; Bergan, Ben-Shaul, and Dulac 2014; Carvalho et al. 2015; Choi et al. 2005). Our previous work supported the theory that the MeApv specifically plays a crucial role in discerning status-related cues early in the olfactory information processing stream, as it responded robustly to dominant cues regardless of familiarity with the individual (W. Lee et al. 2021). This context strengthens our current finding that Glut 8, as one of the few neuronal clusters in the amygdala that shows differential responses, aligns with the MeApv’s specific function.

Interestingly, we observed an opposite response pattern in Glut 0, which was enriched for MeA-CoA-BMA *Ccdc42* neurons. Glut 0 exhibited the highest and second-highest levels of *Oxt* and *Avp* preprohormone expression among all neuronal clusters, respectively. Numerous studies have confirmed the critical role of the oxytocin receptor (OxtR) in the medial amygdala (MeA) for processing social odors and recognizing socially relevant cues (Li et al. 2017; S. Yao et al. 2017). Oxytocin (Oxt) release in the MeA is also implicated in social memory formation (Ferguson et al. 2001). In contrast, the function of vasopressin (Avp) release in the MeA is not as well understood. Similar to OxtR, Avp receptors (Avpr1a and Avpr1b) are essential for social recognition (Albers 2012; Wang et al. 2013) and facilitate social approach behaviours in the MeA (Arakawa, Arakawa, and Deak 2010). Moreover, Avp signaling in the amygdala is linked to aggression promotion (Bosch and Neumann 2010) and exhibits sexually dimorphic expression influenced by androgens, which may underpin its role in aggression (Tong, Abdulai-Saiku, and Vyas 2021). It is important to note that neuropeptide receptor transcripts appeared at low levels in our dataset, which is expected because many of these mRNAs are enriched in dendrites rather than in the nucleus (Cajigas et al., 2012). Since snRNA-seq isolates nuclear RNA only, it tends to underdetect such cytoplasm-localized transcripts (Bakken et al., 2018). Consistent with this, whole-cell scRNA-seq studies of the amygdala report higher receptor detection (O’Leary et al., 2020; Hochgerner et al., 2023).

Glut 0 was also one of two glutamatergic clusters that exhibited low VGLUT1 expression and more prominent VGLUT2 expression compared to other clusters. Indeed, VGLUT2 neurons are the primary glutamatergic subtype intermingled with GABAergic neurons in the MeA and biomedical amygdala (BMA), further validating this cluster’s MMC annotation (Hochgerner et al. 2023). Along with the observed increase in Glut 8 activity in response to dominant cues, these results imply that VGLUT1 and VGLUT2 expressing neurons in the amygdala may have distinct roles in processing social olfactory signals. One of the key distinctions between VGLUT1 and VGLUT2 transporters lies in their association with different glutamate release patterns. VGLUT1 is associated with a low probability of glutamate release, which is linked to a susceptibility to long-term potentiation (LTP), a process that strengthens synaptic connections and supporting learning and memory; in contrast, VGLUT2 is associated with to a high probability of glutamate release and is receptive to long-term depression (LTD), a mechanism that weakens synaptic connections and is also important for the refinement of learning and memory circuits. (Fremeau et al. 2004; Park et al. 2019). How these opposing yet complementary mechanisms underlie differential responses to social stimuli remains unclear. One possible explanation is that activation of VGLUT1-positive neurons in response to dominant cues may enhance the memory of these cues, reinforcing the importance of remembering dominant individuals. In contrast, VGLUT2-positive neuron activation in response to subordinate cues may engage LTD-related mechanisms associated with adaptive forgetting, in which weakening synaptic connections serves to selectively reduce the salience of less behaviorally relevant social information and maintain flexibility in ongoing social interactions (Ryan and Frankland 2022). Further research is required to clarify the unique contributions and potential interactions of these spatially proximal but functionally distinct neuronal populations in processing social cues.

We also observed an increase in response to dominant stimuli in the Glut 9 cluster, which exhibited an expression profile similar to EPd neurons. This observation is consistent with recent studies showing glutamatergic EPd neurons exhibit heightened immediate early gene (IEG) expression following exposure to socially dominant individuals, compared to subordinate ones (Dwortz and Curley 2025). The EPd appears to be multifunctional, interfacing with areas involved in olfaction (e.g., olfactory bulb, medial amygdala, perirhinal cortex), emotional responses (BLA), memory (e.g., entorhinal cortex), and sensorimotor functions (Neafsey, Hurley-Gius, and Arvanitis 1986; Behan and Haberly 1999; Meis et al. 2008; Watson, Smith, and Alloway 2017). Emerging research also suggests that the EPd regulates interoceptive states, given its consistently high baseline activity during exploratory behaviour in mice and during slow-wave sleep in humans (Manjila et al. 2024; Ponomarenko, Korotkova, and Haas 2003). Additionally, some have proposed that EPd activation may contribute to conscious odor perception, as evidenced by situations in which human subjects not only detect an odor but also consciously acknowledge its presence (Traub and Whittington 2022). These findings collectively indicate that the EPd may be involved in integrating internal and external states. Thus, differential EPd activation in response to olfactory cues may indicate the recognition of relative dominance between the subject and the cue.

We also observed differential activity in specific GABAergic neurons. We identified a GABAergic cluster characterized by elevated expression of *Drd2*, which closely resembled striatal DRD2-expressing neurons. This raises the possibility that our dataset includes dorsal striatal neurons in proximity to the CEA. Alternatively, this cluster could consist of striatal-*like* DRD2-expressing GABAergic neurons identified within the CEA itself (McCullough et al. 2018; McDonald 1982). The significance of DRD2 signaling within the amygdala includes its role in modulating sociability (Ike et al. 2024), risk avoidance and impulsive behaviours (Kim et al. 2018; Casey et al. 2022). Thus, DRD2 activation in response to dominant odors may reflect social vigilance and avoidance of potentially agonistic interactions. Furthermore, polymorphisms in the *Drd2* gene are linked to variations in amygdala activity and emotional responses to visual social cues in humans (Blasi et al. 2009). Given these insights, our findings suggest that activation of amygdala *Drd2* neurons is involved in evaluating socially relevant cues across various sensory modalities.

We also performed hdWGCNA analysis on glutamatergic and GABAergic nuclei to investigate the organization of active biological processes in our dataset. This analysis identified modules of co-expressed genes whose expression patterns correlated with specific neuronal clusters, supporting the idea that these clusters represent biologically meaningful groups with specialized roles. Gene Ontology (GO) analysis further illuminated what these active processes might be. For instance, in Glut 9, corresponding to glutamatergic EPd neurons, we observed elevated expression of the green module, which is enriched for genes associated with cell-cell adhesion and junction assembly, suggesting that these processes may be particularly relevant for this population.

Interestingly, not all clusters exhibited overexpressed modules, implying that some clusters might contain neurons serving more generalized functions. For example, Glut 8, associated with glutamatergic MeA neurons, lacked distinctive module overexpression, indicating a potentially broader functional role. Collectively, these findings underscore the cellular diversity in the amygdala, comprising both highly specialized neuronal populations with distinct roles and more generalized populations that may support a range of processes and adapt flexibly to various contexts.

## Conclusion

In conclusion, this study explores how the brain, specifically the amygdala, responds to social olfactory stimuli using snRNA-seq. We have demonstrated distinct activation patterns in specific neuronal populations. Specifically, two groups of VGLUT1 neurons, with transcriptional profiles aligning with the MeA and EPd, respectively, respond more to dominant cues. Conversely, VGLUT2 neurons corresponding to the MeA-Coa-BMA region exhibited elevated responses to subordinate cues. The distinct responses of VGLUT1 and VGLUT2 neurons within the MeA region suggest a MeA-centered circuit may play a key role in mediating responses to social cues. Additionally, the increased expression of both *Avp* and *Oxt* preprohormone mRNAs in the MeA-Coa-BMA population underscores the importance of neuropeptide signaling in regulating social perception. Our results also show that dopamine DRD2 signaling in inhibitory neuron populations is also involved in processing social cues. Future studies should expand the number of sequencing samples to replicate our findings. Moreover, with recent advances reducing costs and increasing the throughput of nuclei analysis (Gezelius et al. 2024), larger and more comprehensive studies are now feasible. Further research should not only confirm these transcriptional responses but also delve into the causal relationships between identified neurotransmitters and amygdala neuronal behaviour, paving the way for targeted interventions to modulate specific neural circuits within the amygdala.

## Acknowledgements

We thank members of the Curley and Hofmann labs for their helpful discussions throughout this work. JPC was supported by the NIH and HAH was supported by the Reeder Endowed Fellowship in Evolutionary Biology. We also thank the Genomic Sequencing and Analysis Facility in the UT Austin Center for Biomedical Research Support (RRID# SCR_021713) for guidance on sample submission as well as library preparation and sequencing. Computational analyses were performed using the Biomedical Research Computing Facility at UT Austin, Center for Biomedical Research Support (RRID#: SCR_021979).

## Data Availability Statement

All data discussed in this article are included in the supplementary materials. Raw sequencing files and processed gene-count matrices are publicly available through the Gene Expression Omnibus (GEO) under accession number GSE310656.

## Supplemental Materials

**Fig S1.**
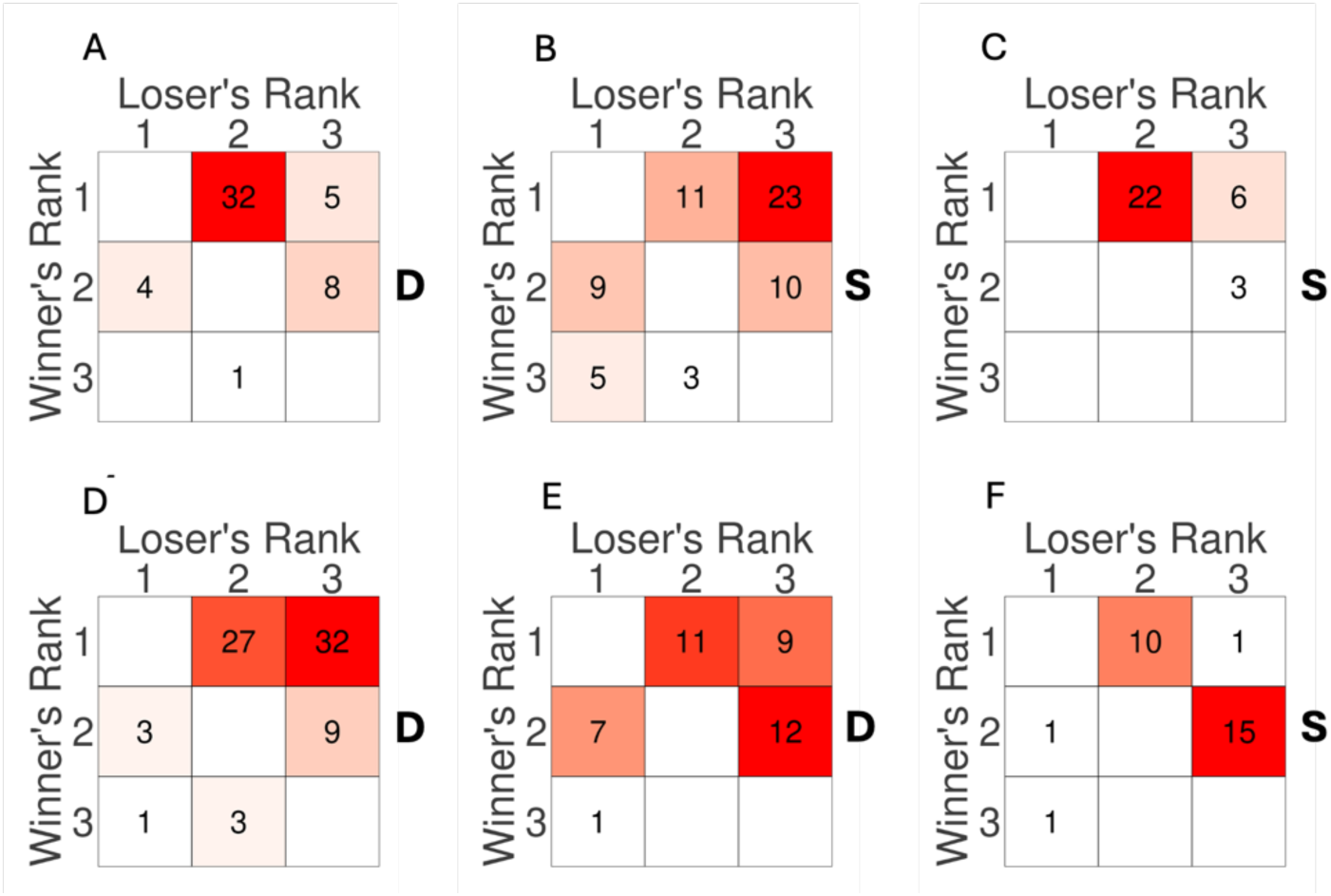
Agonistic Behavior Observations. Sociomatrices show the total frequency of agonistic interactions that occurred between all pairs of individuals across all cohorts (A-F) over the entire observation period. Winners of each contest are listed in rows, and losers are listed in columns. Ranks were calculated using the DS method (see methods). nuclei of each matrix are colored on a gradient from white (lowest value in each matrix) to red (highest value in each matrix). The stimulus presented to each mid-ranked subject is denoted by “D” or “S” for dominant and subordinate urine, respectively.

**Fig S2.**
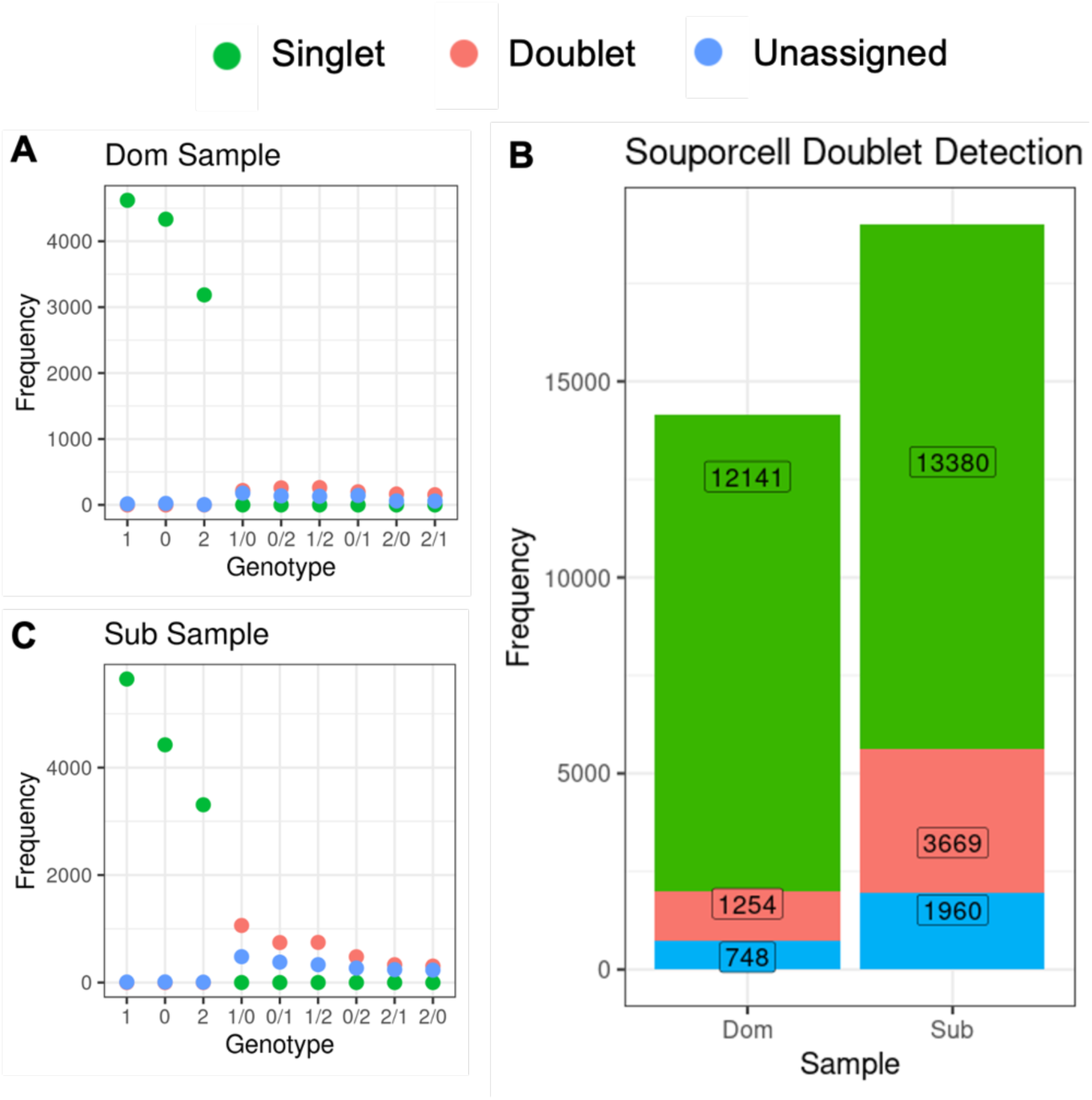
Breakdown of Individual Genotypes per Sample Pool. Results of Souporcell sample demultiplexing. Single nuclei are represented in green. Suspected doublets are in red and nuclei for which the method could not confirm singlet or doublet status are in blue.

**Fig S3.**
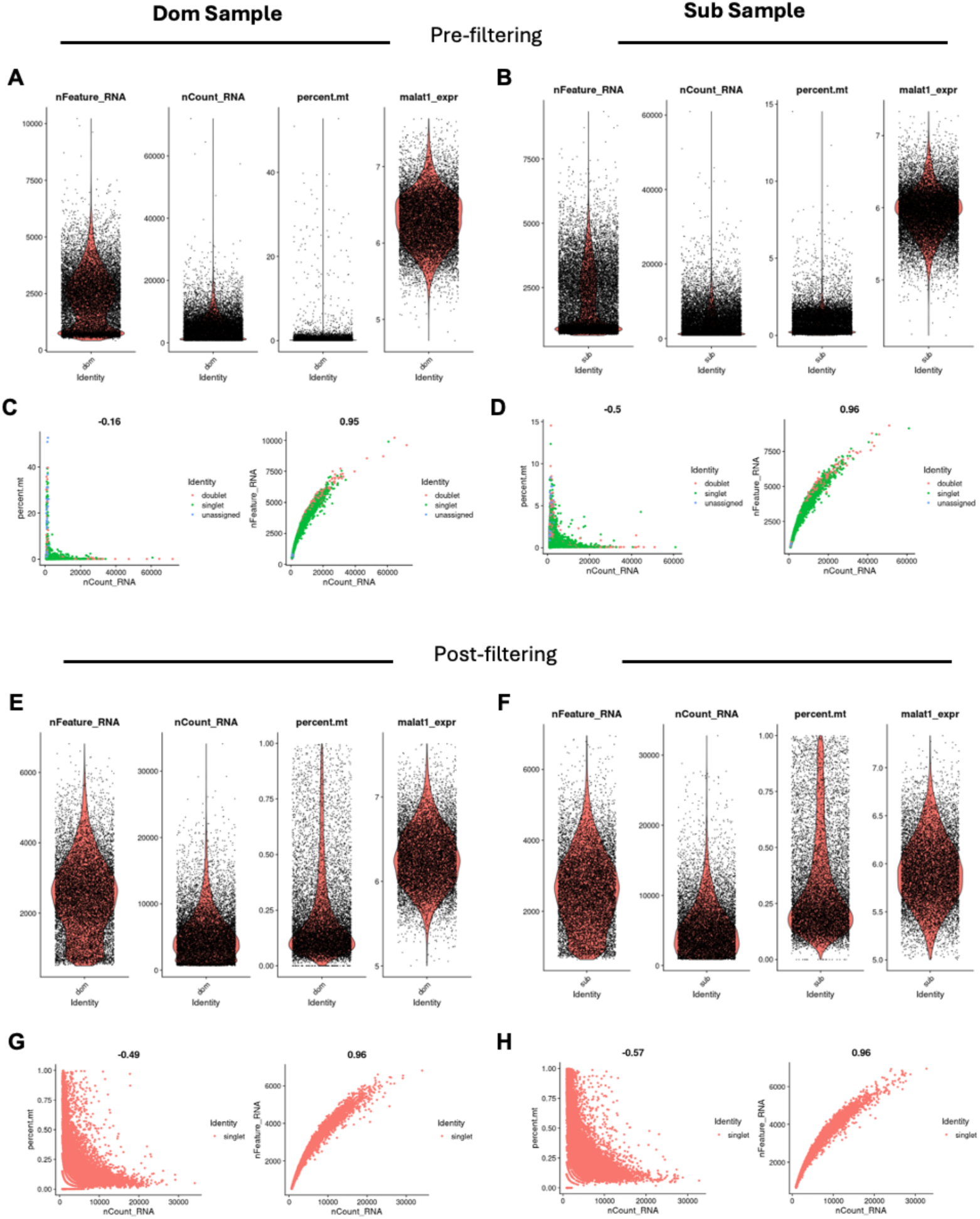
Quality Control Metrics for snRNA-Seq Data Using Seurat. Plots display various quality metrics evaluated for single-cell RNA sequencing data prior to cell filtering (A-D) and after cell filtering (E-H). The left column of plots (A,C,E,G) shows nuclei from the Dom sample, while the right column (B,D,F,H) shows nuclei from the Sub sample. A,B,E, F) Violin plots for quality metrics: nFeature_RNA refers to the number of unique genes detected per cell, aiding in identifying potential doublets; nCount_RNA refers to the total mRNA molecules counted per cell, where extremes may indicate either poor-quality samples or doublets; percent.mt indicates the percentage of mitochondrial RNA content per cell, with higher values typically signifying cell stress or apoptosis; malat1_expr is the expression level of the non-coding RNA MALAT1, useful for assessing cell state or technical variations. Each plot includes a density overlay that highlights the distribution of data points, with individual nuclei represented as black dots. These plots are instrumental in setting thresholds for data filtering. C,D,GH) The left scatter plot shows the relationship between mitochondrial RNA content and total mRNA counts nuclei with low total RNA counts and higher mitochondrial content may be poor quality. The right scatter plot shows the relationship between the number of detected genes and total mRNA counts. Unusually high mRNA counts and a higher number of detected genes may be indicative of doublets. Data points represent nuclei and are color coded by Souporcell assignment as a “singlet”, “doublet”, or “unassigned”. Only singlets were retained for further analysis.

**Fig S4.**
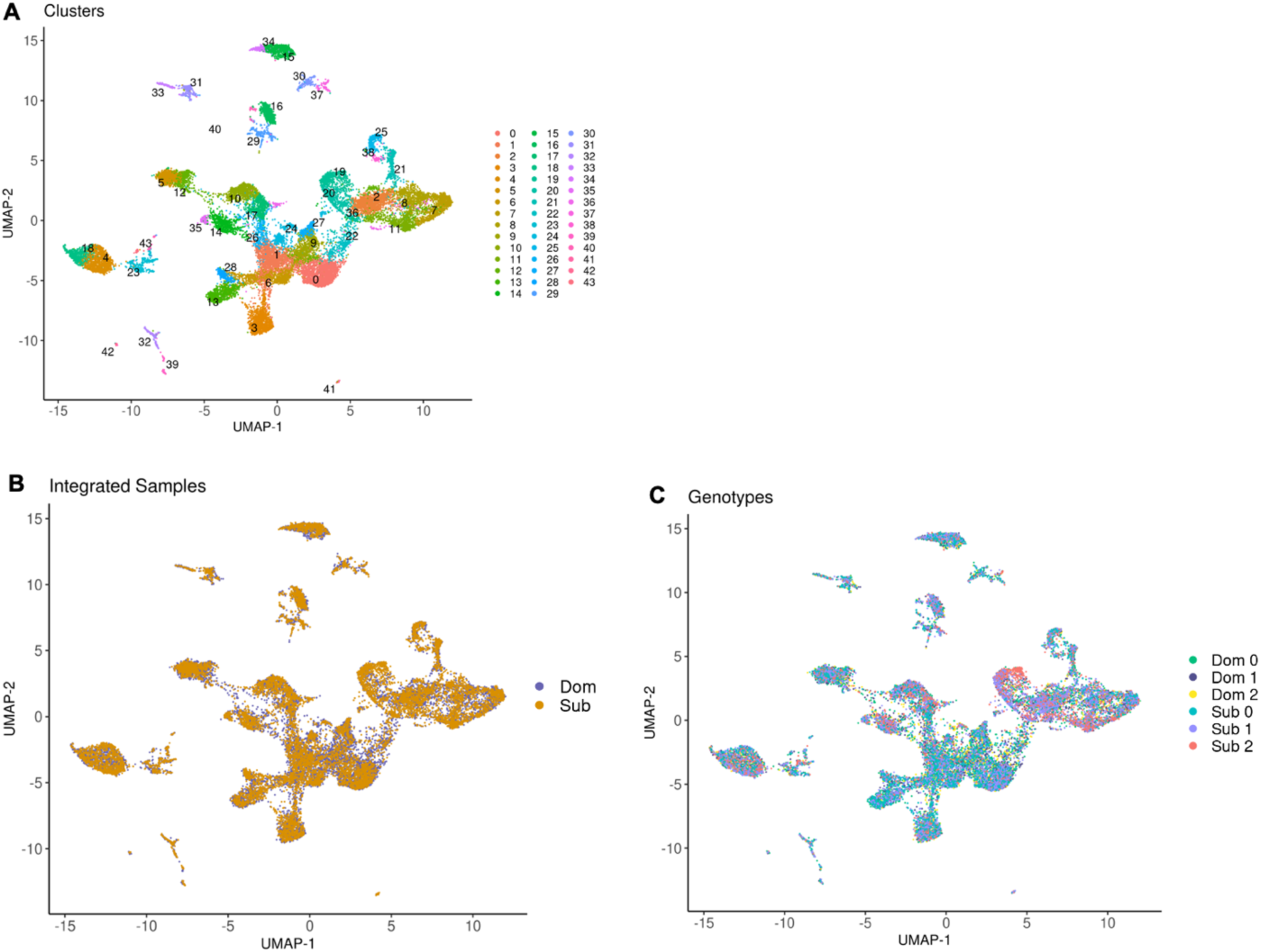
UMAP Visualization of Integrated snRNA-Seq Data from Dominant and Subordinate Samples. A) Uniform Manifold Approximation and Projection (UMAP) plot showing 22,695 nuclei from an integrated dataset combining samples from both dominant (Dom) and subordinate (Sub) groups. The integration process aligns the datasets to minimize batch effects, enabling a unified view of cellular heterogeneity. Each point represents a single cell and is colored by clusters, which consist of transcriptionally similar nuclei in the combined dataset. B) The same UMAP with nuclei color-coded by their sample of origin, visualizing the successful integration of the Dom and Sub datasets. C) The same UMAP with nuclei color-coded according to the individual mouse from which they originated, as determined by Souporcell. Each color represents a distinct genetic profile corresponding to a specific mouse.

**Fig S5.**
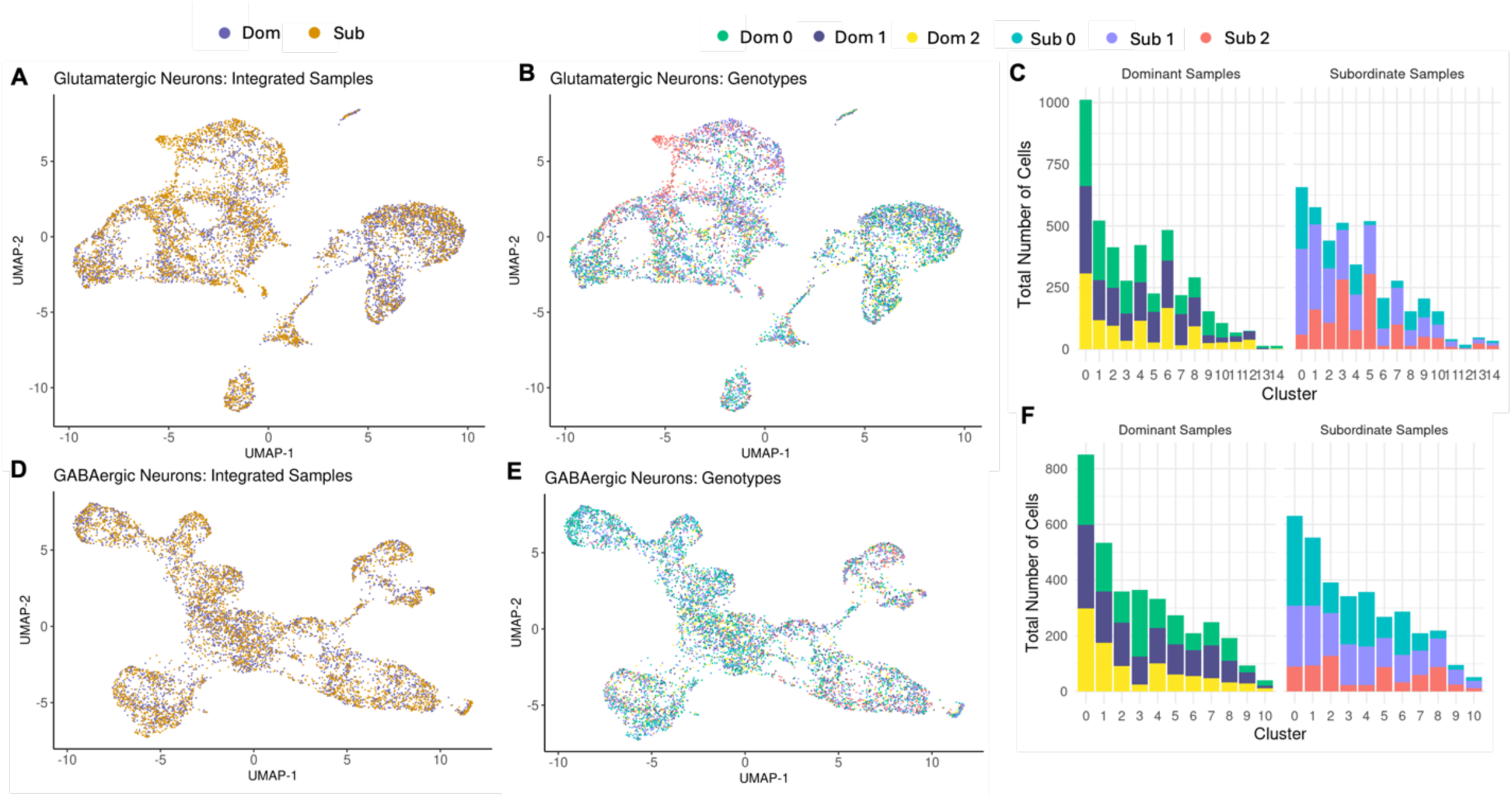
UMAPs and Cluster Distribution of Integrated Glutamatergic and GABAergic Neuron Datasets. A, D) UMAP visualizations of integrated samples for glutamatergic neurons (A) and GABAergic neurons (D), with nuclei colored by their sample origin. B, E) UMAP plots showing the genotypic categorization within the glutamatergic (B) and GABAergic (E) populations. Each color represents a putative individual mouse within dominant (Dom 0, Dom 1, Dom 2) and subordinate (Sub 0, Sub 1, Sub 2) sample pools. C,F) Bar graphs displaying the distribution of nuclei assigned to each genotype across identified clusters within Dom and Sub sample pools for glutamatergic (C) and GABAergic (F) neurons.

**Fig S6.**
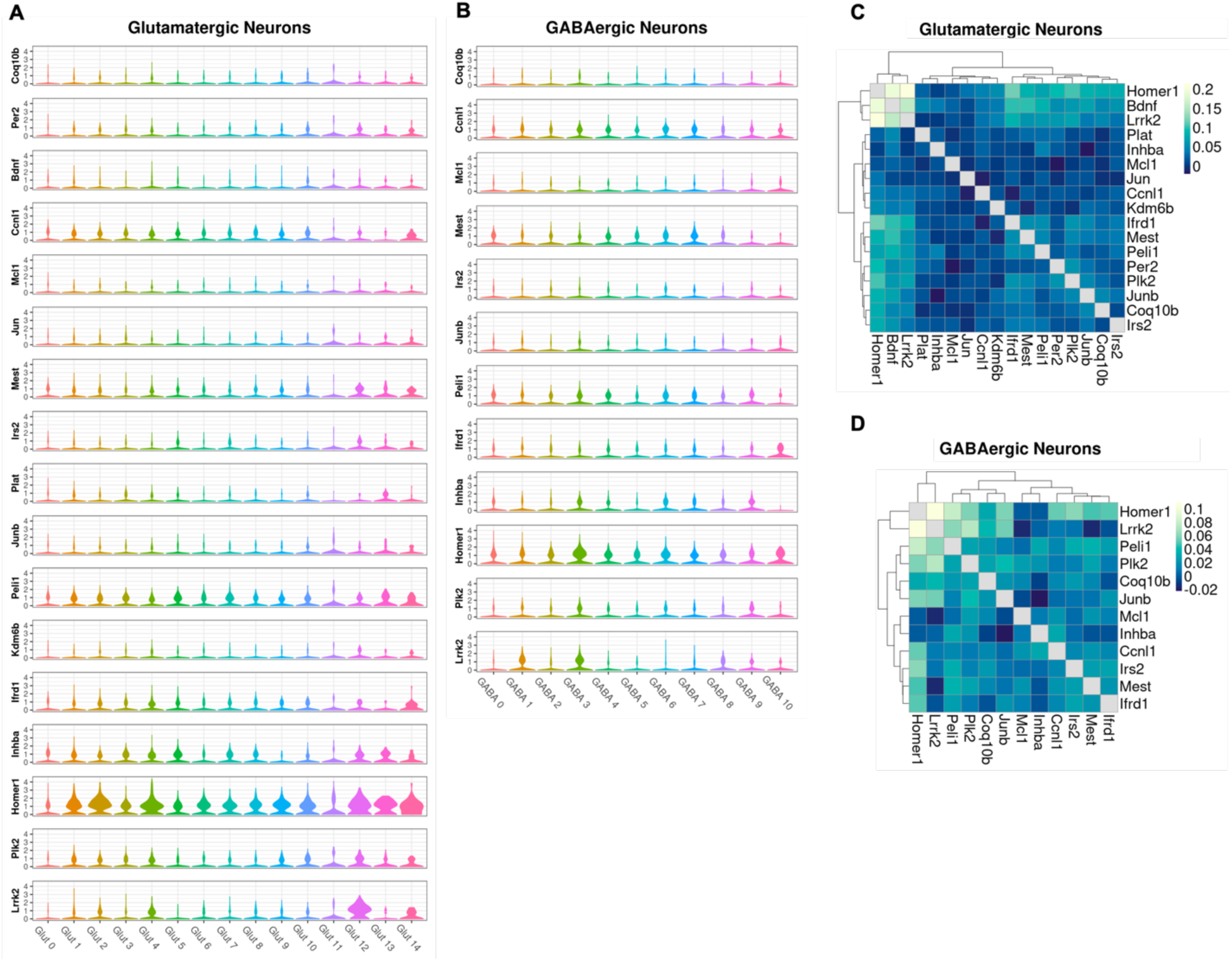
Expression and Correlation Analysis of Immediate Early Genes (IEGs) in Glutamatergic and GABAergic Neurons. A, B) Violin plots showing the log-normalized expression levels of IEGs across glutamatergic (A) and GABAergic (B) neuron clusters. Each plot shows the distribution of expression for a specific gene across different clusters. C, D) Heatmaps depicting the correlation matrices of IEG expression within glutamatergic (C) and GABAergic (D) neurons. Each matrix presents the pairwise Pearson correlation coefficients between the expression levels of IEGs. The color scale, from dark blue to light blue, indicates the strength of the correlation, with lighter shades representing stronger positive correlations.

**Fig S7.**
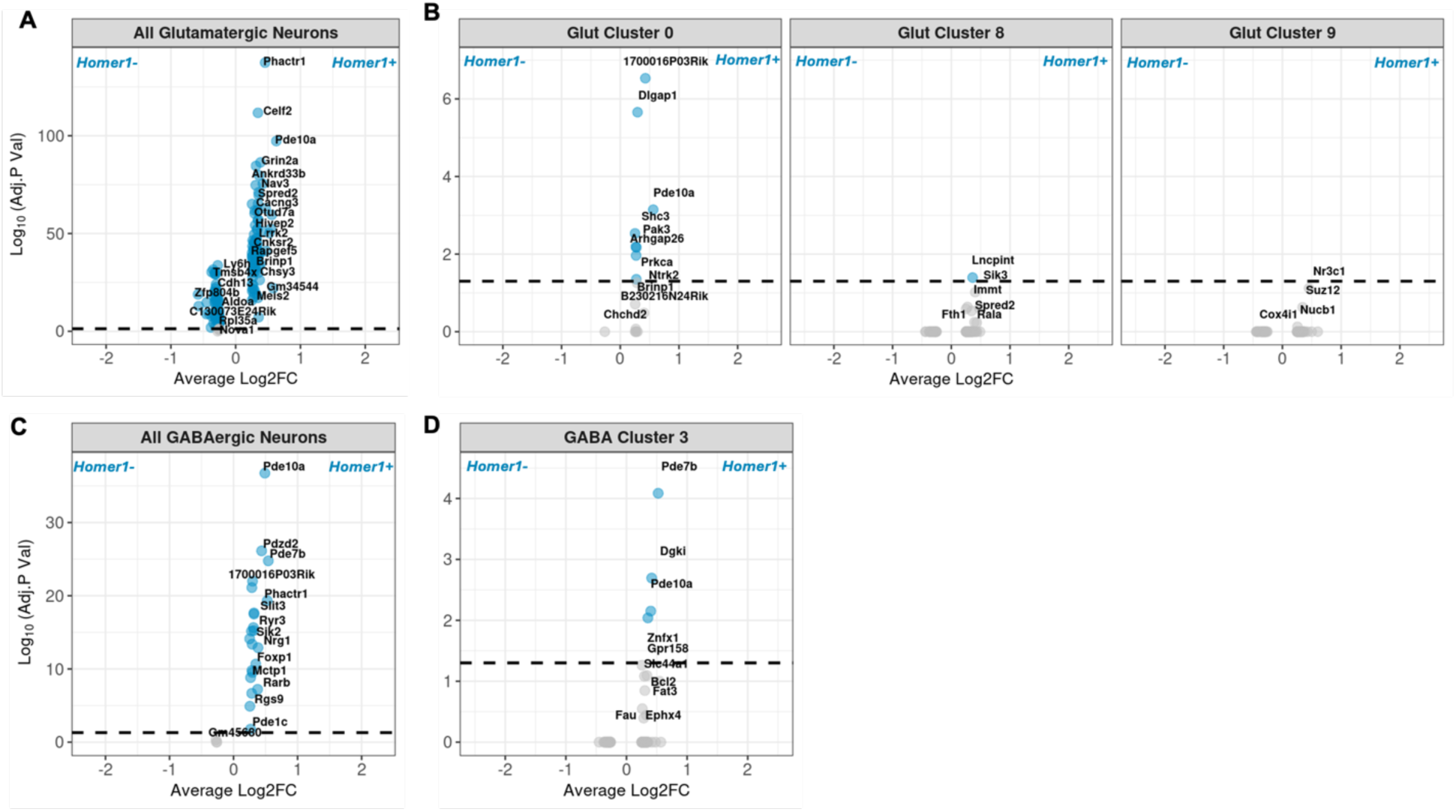
Differentially Expressed Genes between *Homer1⁺* and *Homer1⁻* Nuclei. Volcano plots display the effect size and significance of differentially expressed genes (DEGs) between neurons that express Homer1 (*Homer1⁺*) and those that do not (*Homer1⁻*) among all glutamatergic neurons (A), all GABAergic neurons (C) and clusters that exhibited significant differential proportions of *Homer1⁺*neurons between Dom and Sub stimuli. Data points in blue indicate DEGs with Bonferroni-adjusted P-values of 0.05 or higher. Genes with positive Log2 fold changes (Log2FC) show higher expression in *Homer1⁺* nuclei compared to *Homer1⁻* nuclei. The gene *Homer1* is excluded from these plots because it was used to define the differential expression groups, leading to an infinite value for Log(adjusted P-value).

**Fig S8.**
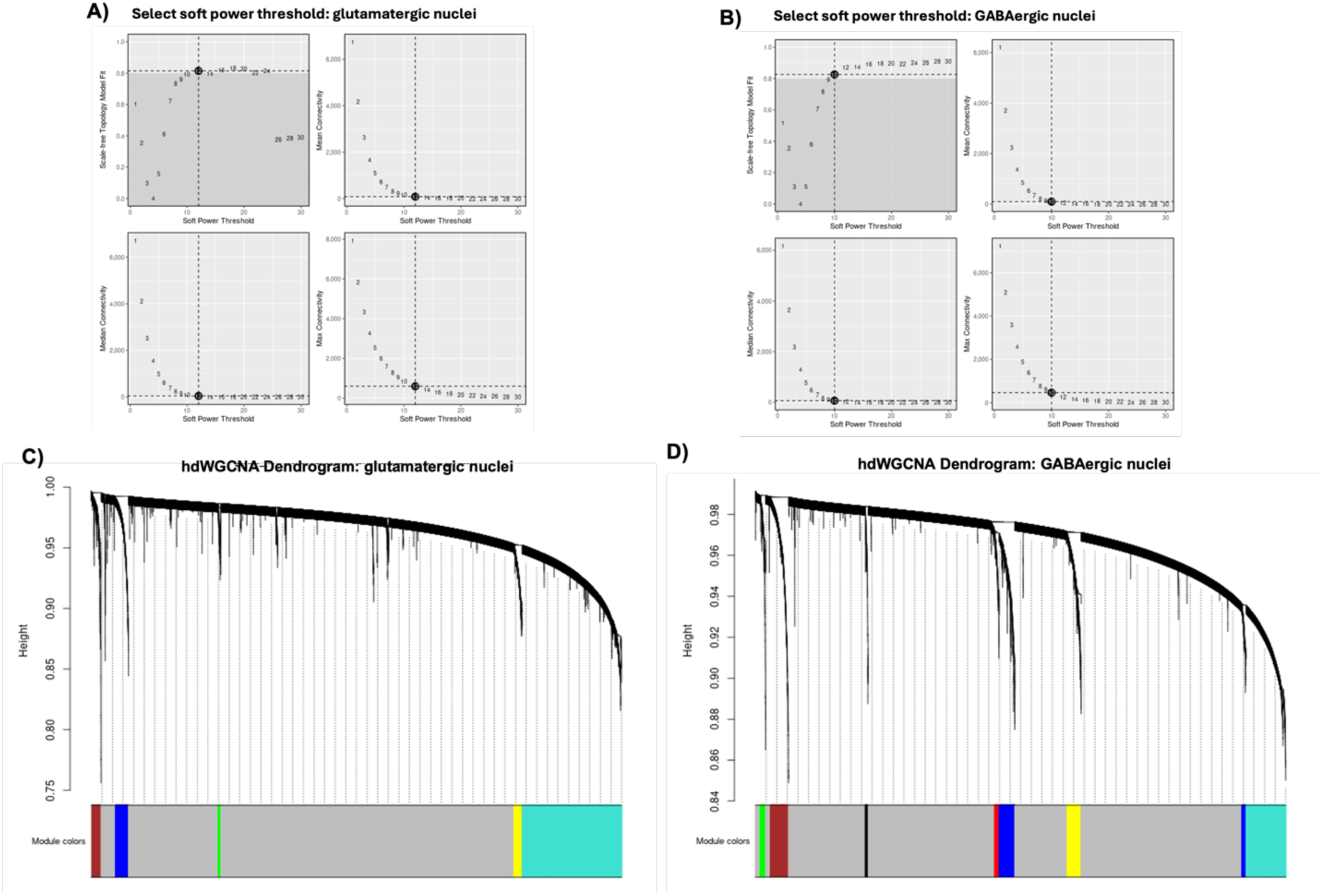
Supplemental hdWGCNA Plots. A,B) Selection of soft power threshold in hdWGCNA for glutamatergic. (A) and GABAergic neurons (B). These plots demonstrate the process of selecting a soft power threshold for network construction in hdWGCNA and were used to determine the scale-free fit index and mean connectivity as functions of the soft power threshold (x-axis). Top Left: Scale-free fit index vs. soft power threshold. The dashed line indicates the target fit index of 0.9. Top Right: Mean connectivity vs. soft power threshold, showing a decrease as the power increases. Bottom Left: A detailed view focusing on lower soft power thresholds and their corresponding scale-free fit indices. Bottom Right: Detailed view of mean connectivity against lower soft power thresholds. The chosen power threshold optimizes network topology for subsequent gene co-expression analysis, balancing between adequate connectivity and a high scale-free fit index. C, D) The dendrogram illustrates the hierarchical clustering of genes based on their topological overlap, used to assess shared connectivity in the hdWGCNA package. Each endpoint of the dendrogram corresponds to a single gene, and the height at which branches merge reflects the dissimilarity between gene expression profiles. The y-axis shows the height (dissimilarity) scale, which quantifies the distance between gene pairs. Lower heights indicate greater similarity. The color band at the bottom denotes the module assignment for each gene. Each color represents a different module, indicating a group of highly interconnected genes. The gray color denotes genes that have not been assigned to any module, often due to insufficient correlation or unique expression patterns that do not align well with other clusters.

**Fig S9.**
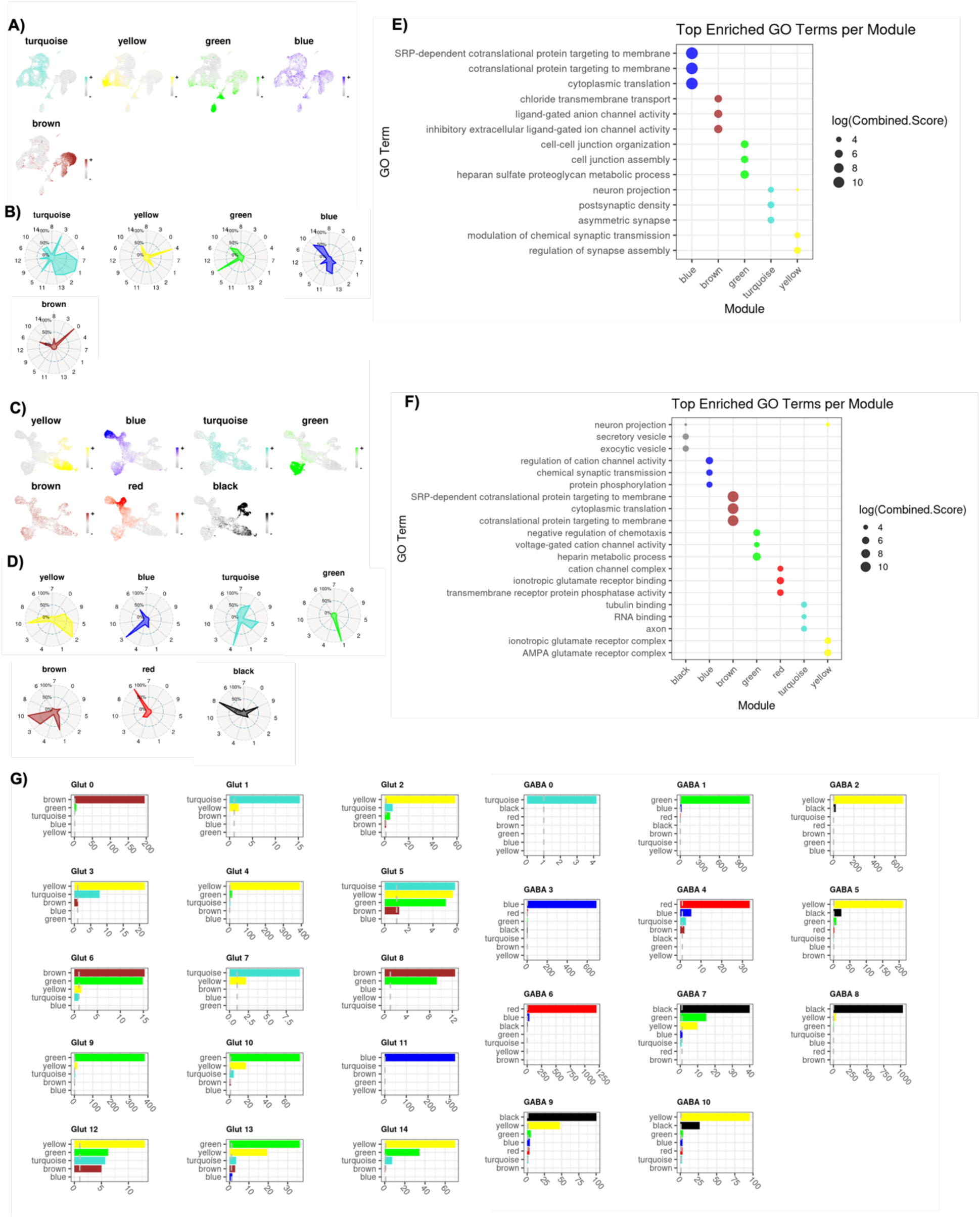
hdWGCNA of Glutamatergic and GABAergic Clusters. A,C) Feature plots showing the expression of module eigengenes (MEs) across UMAP projections of glutamatergic (A) and GABAergic (C) nuclei. The color intensity in each plot corresponds to the expression level of the ME in individual nuclei. B,D) Radar plots depicting the relative expression levels of Mes across the glutamatergic (B) and GABAergic (D) clusters. The radial distance from the center indicates the relative expression level of a given ME within that cluster, with values closer to the outer edge indicating higher expression levels. The center of the circle represents 0% of the maximum expression of the ME, while the outer edge represents 100%. E) F) Dot plots show significant GO terms for each module with the three lowest P values. Dot color corresponds to the module, while size corresponds to the log of the Combined Score for each term. The Combined Score is equal to the log of the P value multiplied by the z-score of enrichment for a given term. G) Bar plots show the number of genes shared between modules and cluster markers. A dashed gray line marks the significance threshold, indicating the minimum level of gene overlap considered statistically significant.

**Table S1.**
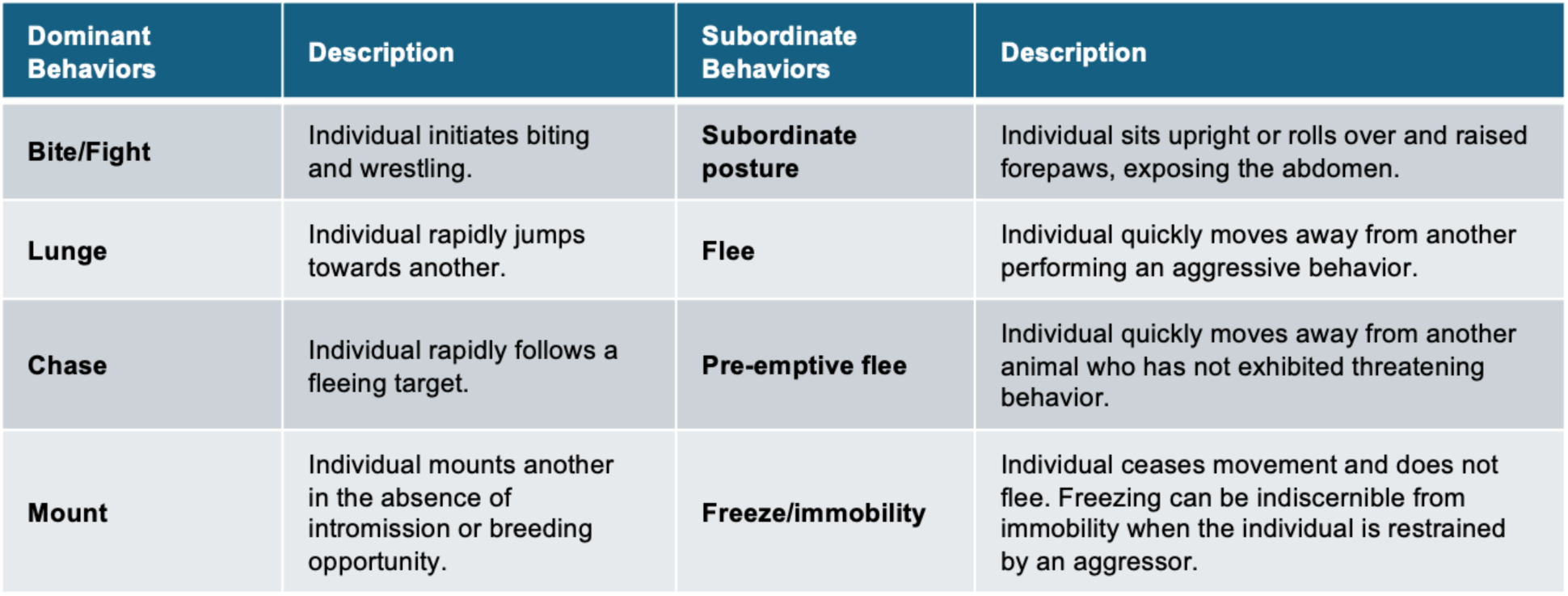
Mouse Social Behavior Ethogram. Observers documented instances of agonistic interactions by recording pairs of offensive and defensive behaviors (e.g., “Bite/Flight – Flee”). In cases of a “pre-emptive flee,” where the aggressive animal displayed no observable behavior, “NA” was recorded.

| Dominant Behaviors | Description | Subordinate Behaviors | Description |
| --- | --- | --- | --- |
| **Bite/Fight** | Individual initiates biting and wrestling. | **Subordinate posture** | Individual sits upright or rolls over and raised forepaws, exposing the abdomen. |
| **Lunge** | Individual rapidly jumps towards another. | **Flee** | Individual quickly moves away from another performing an aggressive behavior. |
| **Chase** | Individual rapidly follows a fleeing target. | **Pre-emptive flee** | Individual quickly moves away from another animal who has not exhibited threatening behavior. |
| **Mount** | Individual mounts another in the absence of intromission or breeding opportunity. | **Freeze/immobility** | Individual ceases movement and does not flee. Freezing can be indiscernible from immobility when the individual is restrained by an aggressor. |

**Table S2.**
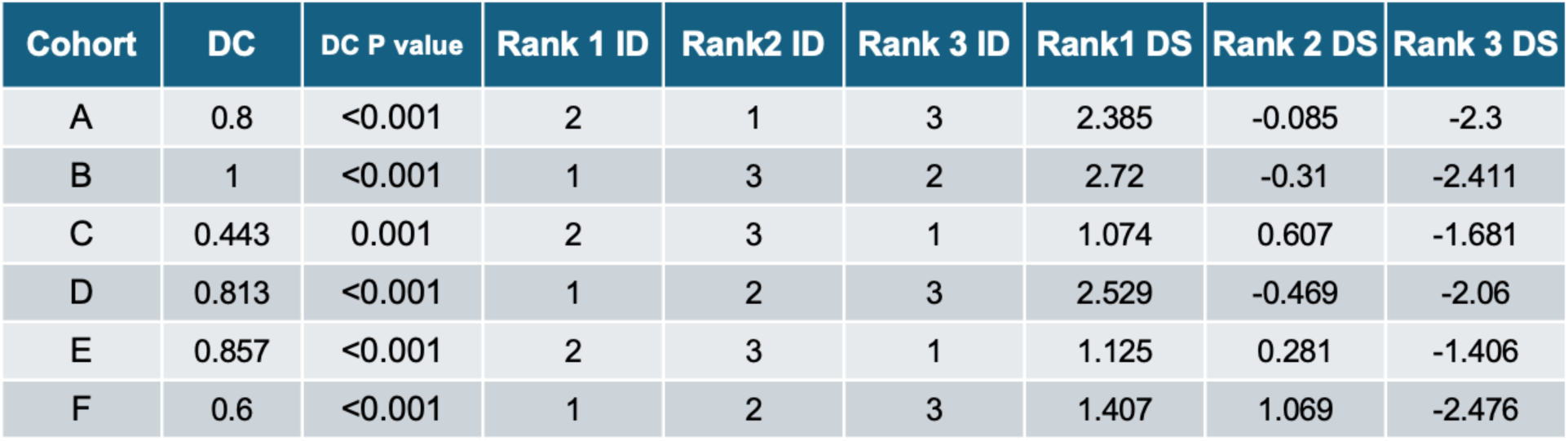
Dominance Results. Directional Consistency (DC) of dominance interactions and David’s Scores (DS) were calculated. In brief, DC assesses the degree to which all agonistic interactions in a group occur in the direction of the more dominant individual to the more subordinate individual within each relationship. It is equal to (H-L)/(H+L) where H is the frequency of behaviors occurring in the most frequent direction and L is the frequency of behaviors occurring in the least frequent direction within each relationship. The significance of DC values were evaluated using the randomization test (Leiva, Solanas, and Salafranca 2008). DS provides an individual dominance rating and ranking for each individual in a group, determining the overall success of each individual at winning contests relative to the success of all other individuals. The most dominant and subordinate individuals in each hierarchy had the highest and lowest DS, respectively.

| Cohort | DC | DC P value | Rank 1 ID | Rank2 ID | Rank 3 ID | Rank1 DS | Rank 2 DS | Rank 3 DS |
| --- | --- | --- | --- | --- | --- | --- | --- | --- |
| A | 0.8 | <0.001 | 2 | 1 | 3 | 2.385 | -0.085 | -2.3 |
| B | 1 | <0.001 | 1 | 3 | 2 | 2.72 | -0.31 | -2.411 |
| C | 0.443 | 0.001 | 2 | 3 | 1 | 1.074 | 0.607 | -1.681 |
| D | 0.813 | <0.001 | 1 | 2 | 3 | 2.529 | -0.469 | -2.06 |
| E | 0.857 | <0.001 | 2 | 3 | 1 | 1.125 | 0.281 | -1.406 |
| F | 0.6 | <0.001 | 1 | 2 | 3 | 1.407 | 1.069 | -2.476 |

**The following large supplemental tables are available in a separate in Supplemental Data file available on GitHub at github.com/mfdwortz/Dwortz_2025_Amygdala_snRNAseq.**

**Table S3.** Cell Type Markers. Significantly higher expressed genes in each main cell type. Genes were considered significant based on a log fold change (logFC) ≥ 1 and p-value ≤ 0.05.

**Table S4.** Glutamatergic Neuron Cluster Markers. Significantly higher expressed genes (cluster markers) for each glutamatergic neuron cluster compared to neurons in other glutamatergic clusters. Identified based on logFC ≥ 1 and p-value ≤ 0.05.

**Table S5.** GABAergic Neuron Cluster Markers. Significantly higher expressed genes (cluster markers) for each GABAergic neuron cluster compared to neurons in other GABAergic clusters. Identified based on logFC ≥ 1 and p-value ≤ 0.05.

**Table S6.** Expression Summary for Neurotransmitter and Receptor Genes of Interest Across Neuronal Clusters. Summary of the expression of genes encoding social neuropeptides, neuromodulators, and their receptors across neuronal clusters. Mean nuclear expression levels and standard deviations are provided for each gene of interest.

**Table S7.** Subclass Assignment for Neuronal Clusters. Subclasses were assigned to cells within each neuronal cluster using the Map My Cells tool from the Allen Institute. Columns include the cell count (number of cells assigned to each subclass), the fraction of cells assigned to the subclass within the respective neuronal type (glutamatergic or GABAergic), and the fraction of cells within each subclass relative to all cells in the corresponding neuronal cluster.

**Table S8.** Binomial GLMM Results. Results of binomial generalized linear mixed models (GLMMs) comparing the proportions of *Homer1⁺* nuclei in response to dominant (Dom) and subordinate (Sub) stimuli across neuronal clusters.

**Table S9.** DEGs Between Homer1⁺ and Homer1⁻ Neurons.

Significantly differentially expressed genes (DEGs) between *Homer1⁺* and *Homer1⁻* neurons. Genes with positive log2 fold-change are elevated in *Homer1⁺*neurons, while genes with negative log2 fold-change are elevated in *Homer1⁻* neurons.

**Table S10.** GO Enrichment Results for DEGs Between *Homer1⁺*and *Homer1⁻*Neurons. Gene Ontology (GO) enrichment analysis results for DEGs between *Homer1⁺* and *Homer1⁻* neurons.

**Table S11.** Glutamatergic Neuron Module Gene Assignments. Module assignments for genes in glutamatergic neurons based on high dimensional weighted gene co-expression network analysis (hdWGCNA). The module membership scores (kME) are shown for each module (turquoise, yellow, green, blue, and brown). The gray module represents genes that were not assigned to any specific co-expression module and are considered unclustered.

**Table S12.** GABAergic Neuron Module Gene Assignments. Module assignments for genes in GABAergic neurons based on high dimensional weighted gene co-expression network analysis (hdWGCNA). The module membership scores (kME) are shown for each module (turquoise, yellow, green, blue, and brown). The gray module represents genes that were not assigned to any specific co-expression module and are considered unclustered.

**Table S13.** GO Enrichment Results for hdWGCNA modules. Gene Ontology (GO) enrichment analysis results for genes in hdWGCNA modules.

## References

1. Adolphs, R. 2001. “The Neurobiology of Social Cognition.” Current Opinion in Neurobiology 11 (2): 231–39. 10.1016/S0959-4388(00)00202-6.

2. Adolphs, Ralph. 2003. “Is the Human Amygdala Specialized for Processing Social Information?” Annals of the New York Academy of Sciences 985 (1): 326–40. 10.1111/j.1749-6632.2003.tb07091.x.

3. Albers, H.E. 2012. “The Regulation of Social Recognition, Social Communication and Aggression: Vasopressin in the Social Behavior Neural Network.” Hormones and Behavior, Oxytocin, Vasopressin and Social Behavior, 61 (3): 283–92. 10.1016/j.yhbeh.2011.10.007.

4. Allen Institute for Brain Science (2011). Allen Reference Atlas – Mouse Brain [brain atlas]. Available from atlas.brain-map.org.

5. Arakawa, H., K. Arakawa, and T. Deak. 2010. “Oxytocin and Vasopressin in the Medial Amygdala Differentially Modulate Approach and Avoidance Behavior toward Illness-Related Social Odor.” Neuroscience 171 (4): 1141–51. 10.1016/j.neuroscience.2010.10.013.

6. Aran, Dvir, A.P. Looney, L. Liu, E. Wu, V. Fong, A. Hsu, S. Chak, et al. 2019. “Reference-Based Analysis of Lung Single-Cell Sequencing Reveals a Transitional Profibrotic Macrophage.” Nature Immunology 20 (2): 163–72. 10.1038/s41590-018-0276-y.

7. Behan, M., and L.B. Haberly. 1999. “Intrinsic and Efferent Connections of the Endopiriform Nucleus in Rat.” Journal of Comparative Neurology 408 (4): 532–48. 10.1002/(SICI)1096-9861(19990614)408:4<532::AID-CNE7>3.0.CO;2-S.

8. Bergan, J.F., Y. Ben-Shaul, and C. Dulac. 2014. “Sex-Specific Processing of Social Cues in the Medial Amygdala.” eLife 3 (June):e02743. 10.7554/eLife.02743.

9. Blasi, G., L.L. Bianco, P. Taurisano, B. Gelao, R. Romano, L. Fazio, A. Papazacharias, et al. 2009. “Functional Variation of the Dopamine D2 Receptor Gene Is Associated with Emotional Control as Well as Brain Activity and Connectivity during Emotion Processing in Humans.” Journal of Neuroscience 29 (47): 14812–19. 10.1523/JNEUROSCI.3609-09.2009.

10. Bosch, O.J., and I.D. Neumann. 2010. “Vasopressin Released within the Central Amygdala Promotes Maternal Aggression.” European Journal of Neuroscience 31 (5): 883–91. 10.1111/j.1460-9568.2010.07115.x.

11. Butler, A., P. Hoffman, P. Smibert, E. Papalexi, and R. Satija. 2018. “Integrating Single-Cell Transcriptomic Data across Different Conditions, Technologies, and Species.” Nature Biotechnology 36 (5): 411–20. 10.1038/nbt.4096.

12. Cajigas, Iván J., Georgi Tushev, Tristan J. Will, Susanne tom Dieck, Nicole Fuerst, and Erin M. Schuman. 2012. “The Local Transcriptome in the Synaptic Neuropil Revealed by Deep Sequencing and High-Resolution Imaging.” Neuron 74 (3): 453–66. 10.1016/j.neuron.2012.02.036.

13. Carvalho, V.M.A., T.S. Nakahara, L. M. Cardozo, M.A.A. Souza, A.P. Camargo, G.Z. Trintinalia, E. Ferraz, and F. Papes. 2015. “Lack of Spatial Segregation in the Representation of Pheromones and Kairomones in the Mouse Medial Amygdala.” Frontiers in Neuroscience 9 (August). 10.3389/fnins.2015.00283.

14. Casey, E., M.E. Avale, A. Kravitz, and M. Rubinstein. 2022. “Partial Ablation of Postsynaptic Dopamine D2 Receptors in the Central Nucleus of the Amygdala Increases Risk Avoidance in Exploratory Tasks.” eNeuro 9 (2): ENEURO.0528-21.2022. 10.1523/ENEURO.0528-21.2022.

15. Chen E., Tan C., Kou Y., Duan Q., Wang Z., Meirelles G., Clark N., Ma’ayan A. Enrichr: interactive and collaborative HTML5 gene list enrichment analysis tool. BMC Bioinformatics. 2013;14:128. doi: 10.1186/1471-2105-14-128.

16. Choi, G.B., H. Dong, A.J. Murphy, D.M. Valenzuela, G.D. Yancopoulos, L.W. Swanson, and D.J. Anderson. 2005. “Lhx6 Delineates a Pathway Mediating Innate Reproductive Behaviors from the Amygdala to the Hypothalamus.” Neuron 46 (4): 647–60. 10.1016/j.neuron.2005.04.011.

17. Clarke, Z.A., and G.D. Bader. 2024. “MALAT1 Expression Indicates Cell Quality in Single-Cell RNA Sequencing Data.” bioRxiv. 10.1101/2024.07.14.603469.

18. Curley, James P. 2016. “Compete: Analyzing Social Hierarchies: R Package Version 0.1.”

19. Dębiec, J. 2005. “Peptides of love and fear: vasopressin and oxytocin modulate the integration of information in the amygdala.” BioEssays 27 (9): 869–73. 10.1002/bies.20301.

20. Dong, H.W., G.D. Petrovich, and L.W. Swanson. 2001. “Topography of Projections from Amygdala to Bed Nuclei of the Stria Terminalis.” Brain Research. Brain Research Reviews 38 (1–2): 192–246. 10.1016/s0165-0173(01)00079-0.

21. Dulac, C. and S. Wagner. 2006. “Genetic Analysis of Brain Circuits Underlying Pheromone Signaling.” Annual Review of Genetics 40 (February):449–67. 10.1146/annurev.genet.39.073003.093937.

22. Dwortz, M.F., J. P. Curley, K. M. Tye, and N. Padilla-Coreano. 2022. “Neural Systems That Facilitate the Representation of Social Rank.” Philosophical Transactions of the Royal Society B: Biological Sciences 377 (1845): 20200444. 10.1098/rstb.2020.0444.

23. Dwortz, M.F., & Curley, J. P. (2025). Capturing dynamic neuronal responses to dominant and subordinate social hierarchy members with catFISH. Neuroscience, 579, 105–119. 10.1016/j.neuroscience.2025.05.025

24. Fadok, J.P., T.M.K. Dickerson, and R.D. Palmiter. 2009. “Dopamine Is Necessary for Cue-Dependent Fear Conditioning.” Journal of Neuroscience 29 (36): 11089–97. 10.1523/JNEUROSCI.1616-09.2009.

25. Ferguson, J.N., J. M. Aldag, T.R. Insel, and L.J. Young. 2001. “Oxytocin in the Medial Amygdala Is Essential for Social Recognition in the Mouse.” The Journal of Neuroscience 21 (20): 8278–85. 10.1523/JNEUROSCI.21-20-08278.2001.

26. Ferretti, V., F. Maltese, G. Contarini, M. Nigro, A. Bonavia, H. Huang, V. Gigliucci, et al. 2019. “Oxytocin Signaling in the Central Amygdala Modulates Emotion Discrimination in Mice.” Current Biology 29 (12): 1938–1953.e6. 10.1016/j.cub.2019.04.070.

27. Fremeau, R.T. Jr., K. Kam, T. Qureshi, J. Johnson, D.R. Copenhagen, J. Storm-Mathisen, F.A. Chaudhry, R.A. Nicoll, and R.H. Edwards. 2004. “Vesicular Glutamate Transporters 1 and 2 Target to Functionally Distinct Synaptic Release Sites.” The Journal of Comparative Neurology 477 (4): 386. 10.1002/cne.20250.

28. Gezelius, H., A.P. Enblad, A. Lundmark, et al. 2024. “Comparison of High-Throughput Single-Cell RNA-Seq Methods for Ex Vivo Drug Screening.” NAR Genomics and Bioinformatics 6 (1): lqae001. 10.1093/nargab/lqae001.

29. Hao, Y., S. Hao, E. Andersen-Nissen, W. M. Mauck, S. Zheng, A. Butler, M. J. Lee, et al. 2021. “Integrated Analysis of Multimodal Single-Cell Data.” Cell 184 (13): 3573–3587.e29. 10.1016/j.cell.2021.04.048.

30. Hao, Y., T. Stuart, M. H. Kowalski, S. Choudhary, P. Hoffman, A. Hartman, A. Srivastava, et al. 2024. “Dictionary Learning for Integrative, Multimodal and Scalable Single-Cell Analysis.” Nature Biotechnology 42 (2): 293–304. 10.1038/s41587-023-01767-y.

31. H., Haynes, A. M. Talman, A. Knights, M. Imaz, D. J. Gaffney, R. Durbin, M. Hemberg, and Mara K. N. Lawniczak. 2020. “Souporcell: Robust Clustering of Single-Cell RNA-Seq Data by Genotype without Reference Genotypes.” Nature Methods 17 (6): 615–20. 10.1038/s41592-020-0820-1.

32. Hochgerner, H., S. Singh, M. Tibi, Z. Lin, N. Skarbianskis, I. Admati, O. Ophir, et al. 2023. “Neuronal Types in the Mouse Amygdala and Their Transcriptional Response to Fear Conditioning.” Nature Neuroscience 26 (12): 2237–49. 10.1038/s41593-023-01469-3.

33. Ianevski, A., A. K. Giri, and T. Aittokallio. 2022. “Fully-Automated and Ultra-Fast Cell-Type Identification Using Specific Marker Combinations from Single-Cell Transcriptomic Data.” Nature Communications 13 (1): 1246. 10.1038/s41467-022-28803-w.

34. Ike, K.G.O., S.J.C. Lamers, S. Kaim, S. F. de Boer, B. Buwalda, J.C. Billeter, and M. J. H. Kas. 2024. “The Human Neuropsychiatric Risk Gene Drd2 Is Necessary for Social Functioning across Evolutionary Distant Species.” Molecular Psychiatry 29 (2): 518–28. 10.1038/s41380-023-02345-z.

35. Janak, P.H., and K.M. Tye. 2015. “From Circuits to Behaviour in the Amygdala.” Nature 517 (7534): 284–92. 10.1038/nature14188.

36. Keshavarzi, S., R.K.P. Sullivan, D. J. Ianno, and P. Sah. 2014. “Functional Properties and Projections of Neurons in the Medial Amygdala.” Journal of Neuroscience 34 (26): 8699–8715. 10.1523/JNEUROSCI.1176-14.2014.

37. Kevetter, G.A., and S.S. Winans. 1981. “Connections of the Corticomedial Amygdala in the Golden Hamster. I. Efferents of the ‘Vomeronasal Amygdala.’” Journal of Comparative Neurology 197 (1): 81–98. 10.1002/cne.901970107.

38. Kim, B., S. Yoon, R. Nakajima, H. J. Lee, H. J. Lim, Y.K. Lee, J.S. Choi, B.J. Yoon, G. J. Augustine, and J.H. Baik. 2018. “Dopamine D2 Receptor-Mediated Circuit from the Central Amygdala to the Bed Nucleus of the Stria Terminalis Regulates Impulsive Behavior.” Proceedings of the National Academy of Sciences of the United States of America 115 (45): E10730–39. 10.1073/pnas.1811664115.

39. Kita, H., and S.T. Kitai. 1990. “Amygdaloid Projections to the Frontal Cortex and the Striatum in the Rat.” Journal of Comparative Neurology 298 (1): 40–49. 10.1002/cne.902980104.

40. Lee, J.H., S. Lee, and J.H. Kim. 2017. “Amygdala Circuits for Fear Memory: A Key Role for Dopamine Regulation.” The Neuroscientist 23 (5): 542–53. 10.1177/1073858416679936.

41. Lee, W., H. N. Dowd, C. Nikain, M. F. Dwortz, E. D. Yang, and James P. Curley. 2021. “Effect of Relative Social Rank within a Social Hierarchy on Neural Activation in Response to Familiar or Unfamiliar Social Signals.” Scientific Reports 11 (1): 2864. 10.1038/s41598-021-82255-8.

42. Li, Y., A. Mathis, B. F. Grewe, J.A. Osterhout, B. Ahanonu, M.J. Schnitzer, V.N. Murthy, and C. Dulac. 2017. “Neuronal Representation of Social Information in the Medial Amygdala of Awake Behaving Mice.” Cell 171 (5): 1176–1190.e17. 10.1016/j.cell.2017.10.015.

43. Manjila, S.B., S. Son, H. Kline, R. Betty, Y. Wu, P. Hyun-Jae, D. Parmaksiz, et al. 2024. “Brain-Wide Connectivity and Novelty Response of the Dorsal Endopiriform Nucleus in Mice.” bioRxiv. 10.1101/2024.09.30.615899.

44. McCullough, K.M., N.P. Daskalakis, G. Gafford, F.G. Morrison, and K.J. Ressler. 2018. “Cell-Type-Specific Interrogation of CeA Drd2 Neurons to Identify Targets for Pharmacological Modulation of Fear Extinction.” Translational Psychiatry 8 (August):164. 10.1038/s41398-018-0190-y.

45. McDonald, A.J. 1982. “Cytoarchitecture of the central amygdaloid nucleus of the rat.” Journal of Comparative Neurology 208 (4): 401–18. 10.1002/cne.902080409.

46. McGarry, L.M., and A.G. Carter. 2017. “Prefrontal Cortex Drives Distinct Projection Neurons in the Basolateral Amygdala.” Cell Reports 21 (6): 1426–33. 10.1016/j.celrep.2017.10.046.

47. Meis, S., J.R. Bergado-Acosta, Y. Yanagawa, K. Obata, O. Stork, and T. Munsch. 2008. “Identification of a Neuropeptide S Responsive Circuitry Shaping Amygdala Activity via the Endopiriform Nucleus.” PLOS ONE 3 (7): e2695. 10.1371/journal.pone.0002695.

48. Mora, M.P. de la, A. Gallegos-Cari, Y. Arizmendi-García, D. Marcellino, and K. Fuxe. 2010. “Role of Dopamine Receptor Mechanisms in the Amygdaloid Modulation of Fear and Anxiety: Structural and Functional Analysis.” *Progress in Neurobiology*, Chemical signaling in the nervous system in health and disease: Nils-Åke Hillarp’s legacy, 90 (2): 198–216. 10.1016/j.pneurobio.2009.10.010.

49. Morabito, S., F. Reese, N. Rahimzadeh, E. Miyoshi, and V. Swarup. 2023. “hdWGCNA Identifies Co-Expression Networks in High-Dimensional Transcriptomics Data.” Cell Reports Methods 3 (6): 100498. 10.1016/j.crmeth.2023.100498.

50. Munuera, J., M. Rigotti, and C.D. Salzman. 2018. “Shared Neural Coding for Social Hierarchy and Reward Value in Primate Amygdala.” Nature Neuroscience 21 (3): 415–23. 10.1038/s41593-018-0082-8.

51. Neafsey, E.J., K. M. Hurley-Gius, and D. Arvanitis. 1986. “The Topographical Organization of Neurons in the Rat Medial Frontal, Insular and Olfactory Cortex Projecting to the Solitary Nucleus, Olfactory Bulb, Periaqueductal Gray and Superior Colliculus.” Brain Research 377 (2): 261–70. 10.1016/0006-8993(86)90867-X.

52. O’Connell, L.A., and H.A. Hofmann. 2011. “The Vertebrate Mesolimbic Reward System and Social Behavior Network: A Comparative Synthesis.” Journal of Comparative Neurology 519 (18): 3599–639. 10.1002/cne.22735.

53. O’Leary, Timothy P, Kaitlin E Sullivan, Lihua Wang, Jody Clements, Andrew L Lemire, and Mark S Cembrowski. 2020. “Extensive and Spatially Variable Within-Cell-Type Heterogeneity across the Basolateral Amygdala.” eLife 9 (September): e59003. 10.7554/eLife.59003.

54. Park, S.K., J.H. Hong, J.K. Jung, H.G. Ko, and Y.C. Bae. 2019. “Vesicular Glutamate Transporter 1 (VGLUT1)– and VGLUT2-Containing Terminals on the Rat Jaw-Closing γ-Motoneurons.” Experimental Neurobiology 28 (4): 451. 10.5607/en.2019.28.4.451.

55. Ponomarenko, A.A., T.M. Korotkova, and H.L. Haas. 2003. “High Frequency (200 Hz) Oscillations and Firing Patterns in the Basolateral Amygdala and Dorsal Endopiriform Nucleus of the Behaving Rat.” Behavioural Brain Research 141 (2): 123–29. 10.1016/s0166-4328(02)00327-3.

56. Raam, T., and W. Hong. 2021. “Organization of Neural Circuits Underlying Social Behavior: A Consideration of the Medial Amygdala.” *Current Opinion in Neurobiology*, The Social Brain, 68 (June):124–36. 10.1016/j.conb.2021.02.008.

57. Rosvold, H.E., A.F. Mirsky, and K.H. Pribram. 1954. “Influence of Amygdalectomy on Social Behavior in Monkeys.” Journal of Comparative and Physiological Psychology 47 (3): 173–78. 10.1037/h0058870.

58. Ryan, T.J., and P.W. Frankland. 2022. “Forgetting as a Form of Adaptive Engram Cell Plasticity.” Nature Reviews Neuroscience 23 (3): 173–86. 10.1038/s41583-021-00548-3.

59. Salem, N.A., L. Manzano, M.W. Keist, O. Ponomareva, A.J. Roberts, M. Roberto, and R. D. Mayfield. 2024. “Cell-Type Brain-Region Specific Changes in Prefrontal Cortex of a Mouse Model of Alcohol Dependence.” Neurobiology of Disease 190 (January):106361. 10.1016/j.nbd.2023.106361.

60. Stuart, T., A. Butler, P. Hoffman, C. Hafemeister, E. Papalexi, W. M. Mauck, Y. Hao, Marlon Stoeckius, P. Smibert, and R. Satija. 2019. “Comprehensive Integration of Single-Cell Data.” Cell 177 (7): 1888–1902.e21. 10.1016/j.cell.2019.05.031.

61. Swanson, L.W., and G.D. Petrovich. 1998. “What Is the Amygdala?” Trends in Neurosciences 21 (8): 323–31. 10.1016/s0166-2236(98)01265-x.

62. Tao, Xu, Steven Finkbeiner, Donald B. Arnold, Adam J. Shaywitz, and Michael E. Greenberg. 1998. “Ca2+ Influx Regulates *BDNF* Transcription by a CREB Family Transcription Factor-Dependent Mechanism.” Neuron 20 (4): 709–26. 10.1016/S0896-6273(00)81010-7.

63. Tong, W.H., S. Abdulai-Saiku, and A. Vyas. 2021. “Arginine Vasopressin in the Medial Amygdala Causes Greater Post-Stress Recruitment of Hypothalamic Vasopressin Neurons.” Molecular Brain 14 (1): 141. 10.1186/s13041-021-00850-2.

64. Traub, R.D., and M.A. Whittington. 2022. “A Hypothesis Concerning Distinct Schemes of Olfactory Activation Evoked by Perceived versus Nonperceived Input.” Proceedings of the National Academy of Sciences 119 (10): e2120093119. 10.1073/pnas.2120093119.

65. Vazdarjanova, A., B. L. McNaughton, C.A. Barnes, P.F. Worley, and J.F. Guzowski. 2002. “Experience-Dependent Coincident Expression of the Effector Immediate-Early Genes *Arc* and *Homer 1a* in Hippocampal and Neocortical Neuronal Networks.” The Journal of Neuroscience 22 (23): 10067–71. 10.1523/JNEUROSCI.22-23-10067.2002.

66. Veyrac, A., G. Wang, M.J. Baum, and J. Bakker. 2011. “The Main and Accessory Olfactory Systems of Female Mice Are Activated Differentially by Dominant versus Subordinate Male Urinary Odors.” Brain Research 1402 (July):20–29. 10.1016/j.brainres.2011.05.035.

67. Vries, H. de. 1995. “An Improved Test of Linearity in Dominance Hierarchies Containing Unknown or Tied Relationships.” Animal Behaviour 50 (5): 1375–89. 10.1016/0003-3472(95)80053-0.

68. Wang, Y., X. Li, L. Li, and Q. Fu. 2013. “Vasopressin in The Medial Amygdala Is Essential For Social Recognition in The Mouse. | EBSCOhost.” March 1, 2013. https://openurl.ebsco.com/contentitem/gcd:86260643?sid=ebsco:plink:crawler&id=ebsco:gcd:86260643.

69. Watson, G.D.R., J.B. Smith, and K. D. Alloway. 2017. “Interhemispheric Connections between the Infralimbic and Entorhinal Cortices: The Endopiriform Nucleus Has Limbic Connections That Parallel the Sensory and Motor Connections of the Claustrum.” The Journal of Comparative Neurology 525 (6): 1363–80. 10.1002/cne.23981.

70. Xiao, B., J.C. Tu, R.S. Petralia, J. P. Yuan, A. Doan, C.D. Breder, A. Ruggiero, A. A. Lanahan, R.J. Wenthold, and P. F. Worley. 1998. “Homer Regulates the Association of Group 1 Metabotropic Glutamate Receptors with Multivalent Complexes of Homer-Related, Synaptic Proteins.” Neuron 21 (4): 707–16. 10.1016/S0896-6273(00)80588-7.

71. Yao, S., J. Bergan, A. Lanjuin, and C. Dulac. 2017. “Oxytocin Signaling in the Medial Amygdala Is Required for Sex Discrimination of Social Cues.” Edited by Richard D Palmiter. eLife 6 (December):e31373. 10.7554/eLife.31373.

72. Yao, Z., C.T.J. van Velthoven, M. Kunst, M. Zhang, D. McMillen, C. Lee, W. Jung, et al. 2023. “A High-Resolution Transcriptomic and Spatial Atlas of Cell Types in the Whole Mouse Brain.” Nature 624 (7991): 317–32. 10.1038/s41586-023-06812-z.

73. Yu, Bin, Q. Zhang, L. Lin, X. Zhou, W. Ma, S. Wen, C. Li, et al. 2023. “Molecular and Cellular Evolution of the Amygdala across Species Analyzed by Single-Nucleus Transcriptome Profiling.” Cell Discovery 9 (February):19. 10.1038/s41421-022-00506-y.

74. Zink, C.F., Y. Tong, Q. Chen, D. S. Bassett, J. L. Stein, and A. Meyer-Lindenberg. 2008. “Know Your Place: Neural Processing of Social Hierarchy in Humans.” Neuron 58 (2): 273–83. 10.1016/j.neuron.2008.01.025.

